# Initial TCR Signal Strength Imprints GATA3 and Tbet Expression Driving T-helper Cell Fate Decisions

**DOI:** 10.1101/2024.07.18.604067

**Authors:** Mohammad Ameen Al-Aghbar, Taushif Khan, Meritxell Espino Guarch, Douglas C Palmer, Nicholas van Panhuys

**Affiliations:** Laboratory of Immunoregulation, Human Immunology Division, Sidra Medicine, Doha, Qatar; Jackson Laboratory, Bar Harbor, MN, USA; Surgery Branch, National Cancer Institute (NCI), National Institutes of Health, Bethesda, MD, USA; Lymphocyte Biology Section, Laboratory of Immune System Biology, National Institute of Allergy and Infectious Diseases, National Institutes of Health, Bethesda, MD, USA

## Abstract

With the exception of the T-helper 2 (Th2) subset, cytokine driven pathways provide a robust mechanistic explanation for the observed outcomes of CD4+ T-cell differentiation. Using a quantitative model of activation, we studied the integration of TCR-signal-strength with cytokine signalling during Th2 differentiation. Upon the initial activation of Th-naïve cells, TCR signalling was found to set early expression levels for the master regulators of differentiation Tbet and GATA3, independent of the presence of polarizing cytokines.

Subsequently cytokine stimuli modulated transcription factor (TF) expression levels to tune the outcome of differentiation. Here, weak TCR signalling was sufficient to drive the early upregulation of GATA3 and induce Th2 differentiation, in an IL-4 independent manner. Th1 differentiation was however shown to require additional cytokine signalling input, either in the form of autocrine IFNγ or exogenous IL-12. Using mathematical modelling we demonstrate that T-helper differentiation occurs along a continuum of states. Set by the relative co-expression of regulatory TFs, where effector cytokine production is controlled in a probabilistic manner determined by the relative levels GATA3 and Tbet expressed.

Together, our data indicate TCR signalling inputs drive an early bifurcation in the T-helper differentiation pathway. Together, the integration of TCR signal strength with cytokine inputs act as a mechanism for the detection of immuno-evasive parasitic infections, whilst providing an additional checkpoint to prevent aberrant Th1 associated immunopathology.

## Introduction

Classically T-helper cell differentiation has been described as being driven by qualitative cytokine inputs, providing the third signal in the activation/differentiation cascade, following TCR (signal 1)- and CD28/costimulatory molecule (signal 2)-induced activation. Where, IL-12 [1] and/or IFNγ [2] drive Th1 differentiation, IL-4 induces Th2 differentiation [3], TGFβ/IL-10 polarize regulatory T (Treg) cells and TGFβ/IL-6/IL-23 determines Th17 differentiation[4]. The induction of Th1 differentiation from naïve CD4+ T (Th-naive) cells by dendritic cells (DCs) has been well elucidated, with the discovery of pathogen-associated molecular patterns (PAMPs) and the signalling pathways that allow DCs to transduce the signals generated by specific bacterial and viral agents. In turn, leading to the production of differentiation-inducing cytokines, which provide signal 3 for Th1 differentiation [5].

However, identifying the key signalling factors that induce Th2 differentiation has proved more elusive, with several studies demonstrating IL-4 to be dispensable for *in vivo* Th2 differentiation [6; 7; 8; 9; 10]. The cytokines generated by basophils, epithelial cells, and innate leukocytes (ILC2) which are associated with type II inflammation, including IL-25, IL- 33, and TSLP, have been shown to have significant roles in the generation and characteristics of the Th2 response[11]. However, these factors do not appear to account for the initial polarization of Th-naïve cells towards the Th2 state under *in vivo* conditions [12], and currently no definitive signal 3 for Th2 differentiation has been identified.[13]. As such, it has been theorized that Th2 differentiation may be the result of a default or endogenous pathway[14] [15]. Type-2 conventional DCs (cDC2) have been identified as the *in vivo* initiators of Th2 differentiation [16; 17], where the induction of an IRF4[18] and KLF4[17; 19] driven program in cDC2 results in Th2 differentiation. As cDC2 do not provide an initial source of IL-4, this has furthered the Th2 paradox [20; 21; 22]. Leading to the hypothesis that Th2 differentiation may be an exception to the three-signal paradigm and may occur occur through repression of TLRs, inhibition of TLR signalling and down regulation of signal 3 cytokine production. [13].

A significant body of work has also demonstrated that quantitative signals imparted through the TCR during activation of Th-naïve cells is a key driver of differentiation; with strong TCR stimuli leading to the induction of Th1 differentiation and weak stimuli inducing Th2 differentiation [23; 24; 25; 26; 27]. Previously [28], we demonstrated TCR-driven signalling could direct Th-naive differentiation under *in vivo* conditions. Here, weak TCR signalling was associated with shorter interactions between Th-naive cells and DCs during activation, whereas strong TCR signalling induced a rapid transition to the formation of stable immune synapse-associated interactions. Additionally, it has been shown that the cytokines and their cognate receptors which induce Th1 differentiation are localized to the immune synapse during activation [29]. Whereas IL-4R is not specifically recruited to the immune synapse [30], and Th2 associated cytokines such as IL-2 and IL-4 are thought to be secreted in a non-directional manner [31], indicating that stabilized long-term interactions may be required for Th1 differentiation.

Many of the pathways and programs associated with the ability of Th-naive cells to differentiate into mature T-helper cell subsets have been well elucidated [4]. However, significant discrepancies remain unaccounted for in our understanding of how signals generated by TCR-associated activation integrate with cytokine signalling. In this study, we used a quantitative model of Th-naive activation [27] to assess the precise involvement of both TCR signal strength and polarizing cytokines in the process of Th differentiation.

## Methods

### Mice

B10.A mice of RAG2^−/−^ 5C.C7 transgenic TCR specific for pigeon cytochrome C peptide (pPCC, residues 88-104; KAERADLIAYLKQATAK) on I-Ek, 5C.C7 TCR transgenic RAG2^−/−^ IL- 4^G4/G4^, 5C.C7 TCR transgenic RAG2^−/−^ IL-4^G4/G4^, 5C.C7 TCR transgenic RAG2^−/−^ IFNγ^−/−^, and 5C.C7 TCR transgenic RAG2^−/−^ IFNγ^−/−^ IL-4^G4/G4^ were bred and maintained under specific pathogen-free conditions in the National Institute of Allergy and Infectious Diseases Animal Facility or in the Laboratory Animal Research Centre, Qatar University and used at 6-12 weeks of age. All procedures were approved by the NIAID Animal Care and Use Committee or the Qatar University Animal Care and Use Committee.

### Reagents

For *in vitro* culture; RPMI 1640 (Gibco) was supplemented with 10% fetal calf serum (FCS), 2- ME, glutamine, penicillin, streptomycin, and sodium pyruvate (referred to as cRPMI). Pigeon cytochrome C peptide (residues 88-104; KAERADLIAYLKQATAK) was purchased from either American Peptide Company or Anaspec. Murine rIL-12 and rIL-4 were purchased from R&D. For cell staining, CFSE (Invitrogen); Live/Dead fixable violet dead cell stain (Invitrogen); and anti-mouse antibodies APC/Cy7-CD4 (GK1.5, Biolegend), AF647-CD44 (IM7, Biolegend), PE- CD62L (MEL-14, Biolegend, Biotin-CD49b (DX5, Biolegend), streptavidin-AF488 (Biolegend), PE-Cy7-CD69 (H1.2F3, Biolegend), PE/Cy7-IL-4 (11B11, Biolegend), PerCP-Cy5.5-IFNγ (XMGL2, Biolegend), AF488- or PE-GATA3 (TWAJ, Invitrogen), AF647-Tbet (4B10, Biolegend), AF647-Foxp3 (clone 150D, Biolegend), and PE-RORγt (clone Q31-378, BD) were used. For the MHCII blockade experiments, anti-mouse MHCII (14-4-4S) was purchased from eBiosciences. Phorbol myristate acetate (PMA, Sigma), ionomycin (Sigma), monensin (Calbiochem), and eBioscience Foxp3/TF Staining Buffer Set (ThermoFisher) were used for cell reactivation and fixation.

### Isolation and *in vitro* culture of Th-naive

Th-naive cells were isolated from spleens and lymph nodes of animals as indicated. Isolated cells were stained with Live/Dead, anti-CD4, anti-CD62L, anti-CD44, and anti-DX5; CD44-low CD62L-high CD4+ DX5-T cells were sorted by FACS Aria (Becton Dickinson) to >98% purity and used as Th-naive cells. Where indicated, Th-naive cells were labelled with 1.25 µM CFSE (Invitrogen) before use, according to the manufacturer’s instructions. Th-naive (5 x 10^5^) were cultured in cRPMI in a 48-well plate with P13.9 fibroblast cells [32] (1.25 x 10^5^) expressing I-Ek, CD80, and ICAM-1 which had been treated with 25 µg/ml mitomycin C and loaded with 0.01-10 µM pPCC; 0.01-10 ng rIl-12, 10 ng rIL-4, or 20 µg/ml anti-MHCII were added to cell cultures as indicated.

### Intracellular staining of CD4+ T cells

Following 96 h of stimulation activated CD4+ T cells were restimulated in cRPMI with PMA (100 ng/ml) and ionomycin (1 µg/ml) at 37°C for 4 h in the presence of monensin (2 mM). Cells were surface stained with anti-CD4, and live-cell staining was performed with Live/Dead fixable violet dead cell stain. Cells were fixed and permeabilized with eBioscience Foxp3/TF Staining Buffer Set according to the manufacturer’s instructions.

Intracellular cytokine staining was performed with anti-IFNγ and anti-IL-4 antibodies, and TF staining was performed for GATA3, Tbet, Foxp3, or RORγt, as indicated. Flow cytometric data were collected on either an LSR II (BD Biosciences) or an Aurora (Cytek) flow cytometer and analysed using FlowJo software (TreeStar). For flow cytometric analysis, cells were gated on FSC/SSC to remove debris and non-activated cells subsets, FSC-H/FSC-A to remove doublets, and Live/Dead fixable violet dead cell stain-negative and CD4-positive populations prior to analysis to generate data on the proportions (%) of cytokine- and transcription-factor-expressing cells and mean fluorescence intensities (MFIs).

### RNA sequencing and analysis

RNA was extracted from P13.9, Th-naïve and activated CD4+ T cells following 96 h stimulation with pPCC (without restimulation), using TRizol (ThermoFisher), as per the manufacturer’s instructions. RNA quantification and library preparation for Lexogen QuantSeq 3′ mRNA-Seq was carried out as previously published [33]. Lexogen QuantSeq3′ mRNA-sequencing libraries were prepared according to the manufacturer’s instructions. Libraries were pooled and sequenced on the Illumina NextSeq 500 system to a depth of 4 million reads per library. The initial quality of the sequencing was assessed with FASTQC (v.0.11.8). Reads were aligned against the mouse genome mm10 in STAR v2.6.1d., HTSeq-count (v0.9.1) was used to generate raw counts, and genes were median normalized by DESeq2 and are presented as normalized count values[34].

Gene expression analysis was carried out using packages in iDEP.96 DESEQ [35], and genes were log-transformed with log(x+c), where c = 1 and missing values were treated as zero. Gene set enrichment analysis (GSEA) was conducted with GSEA_4.3.2 and carried out as published [36], using previously defined Th1 and Th2 gene sets[37] derived from mouse CD4+ Th subsets [38]. Differentially expressed genes (DEGs) between pairwise comparisons were identified using DESeq2 (FDR < 0.05, fold change >1.5) in iDEP.96. Ciiider software analysis was used to determine the enrichment of TF binding sites (TFBS), as in Gearing, *et al* [39]. Potential TF binding sites across a promoter region spanning 1500bp upstream and 500bp downstream of the transcription start site of the DEGs identified were scanned by CiiiDER, with a deficit cut-off of 0.15 and enrichment coverage p value =0.05. K-means clustering was used to filter genes that were highly expressed in P13.9 cells and the resultant gene list used as input for partial least squares-discriminant analysis (PL-SDA) using the corresponding package in Metaboanalyst[40]. Top 1% positive and negative genes from PLSDA PC2 loading values were extracted and CPM values plotted. Gene ontology analysis of Biological Processes present in positive and negative genes identified was conducted with WebGestalt 2024[41].

### Microscopy and live cell imaging

Cells were imaged with a Zeiss 710 microscope equipped with a Chameleon laser (Coherent) tuned to 800 nm with a 20X water-dipping lens (NA 1.0, Zeiss) and Zen 2010 acquisition software. Imaging was conducted in enclosed environmental chambers kept at 37°C. Cells were incubated in cRPMI containing 25 mM HEPES. To visualize cells, P13.9 antigen-presenting cells (APCs) were labelled with 100 µM 7-amino-4-chloromethylcoumarin (Invitrogen).

For Ca2+ flux assessment, CD4+ T cells were co-stained with 1.25 µM CytoTrace™ Red CMTPX (Invitrogen) and 2.5 µM Fluo-4 (Invitrogen), in the presence of 0.1% Pluronic-F127 (Invitrogen). Calcium flux analysis was conducted as described previously [28].

### Data Modelling

GATA3:Tbet ratios for individual cells were calculated using the ‘derive parameter’ function in FlowJo to calculate ratios on a single-cell basis. The geometric means (GMs) for the specified populations were then extracted and used for modelling data points.

Association between cytokines and TFs were examined using two linear modelling approaches: generalized linear modelling (GLM) and robust linear modelling (RLM). Both cytokines (IFNγ and IL-4) were modelled independently to measure the effects of the TFs (GATA3 and Tbet) and their ratio (GATA3:Tbet). For each model, the designed matrix was created, with cytokines as independent variables and TFs as dependent (or response) variables. We used the Python (v 3.9) and Statsmodels (v. 0.13.5) packages for conducting analyses.

For TF protein quantification, relative units (r.u.) were determined from the GM indices and were calculated as the GM of the expressing population divided by the GM of the total population. Unless indicated otherwise, GM indices of plotted cell subsets were normalized to those of total populations of cells. Normalized fractions of cytokine-expressing cells were determined as a proportion of each of the TF high (Hi), intermediate (Int), and low (Lo) groups and normalized to the cytokine expression of the total cell population for either 10.0 µM or 0.01 µM pPCC-stimulated cells, as indicated in the relevant figures.

An empirical dose-response curve, y = Ymax × (TF/K + TF), as used in Helmstetter et al. [42], was initially used to describe the functional relationship of GATA3, Tbet, or GATA3:Tbet TF (TF) expression and the frequency of cytokine producers (IL-4 or IFNγ) Y. The two parameters Ymax and K were estimated using the TF expression values and the corresponding frequency of Y producers following restimulation.

Data fitting based on the Hill equation was modelled with the following function:

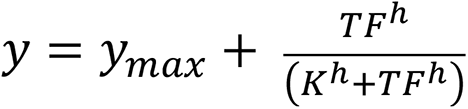

Where y = % cytokine expression (IL-4 or IFNγ) in a given population following restimulation; y_max_ = maximum predicted cytokine expression from the populations; TF = MFI of either GATA3 or Tbet for the population or GATA3:Tbet ratio, as calculated from GM data; K = TF MFI at which %y producers is half-maximal; h = Hill constant; data were fitted for each cytokine (y), initiated with y_max_ with a maximum value of y; h = 0 and K=10, and the curve fitting was minimized on TF and cytokine expression. The final learned values of these parameters were fitted to plot the curves presented.

### Statistical calculations

Data are routinely presented as ±S.E.M. One-way ANOVA with Tukey’s post-testing were used for the statistical analysis of multiple groups. Student’s t test (two-tailed) was used for the statistical analysis of differences between two groups. Data were analysed using Prism Graph-Pad 9 software. P = *,<0.05; **,<0.01; ***<0.001.

## Results

### Quantitative Signalling Alters Expression of GATA3 and Tbet at the Single-Cell Level

As the effects of varying TCR-signal strength on altering the outcome of differentiation have previously been studied through analysis of effector cytokine production [23; 24; 25; 26; 27]. We aimed to determine the effects of varying TCR signalling on the induction of lineage defining TFs at the single-cell level. As in Yamane *et al.* [27], P13.9 fibroblasts expressing MHCII, CD80, and ICAM-1 were used as APCs. Here, P13.9 cells were pre-loaded with peptide by incubation in the presence of pPCC at concentrations of 0.01µM to 10.0µM, allowing them stimulate Th-naïve cells with defined peptide:MHC (p:MHC) levels.

Naïve TCR transgenic 5CC7 Rag^−/−^ CD4+ T cells were activated in the presence of pPCC- loaded APCs and on day 4 following activation cells were restimulated, and GATA3 and Tbet expression examined. In comparison to Th-naive cells, activated cells exhibited a marked upregulation of GATA3 under all conditions (Fig. S1). Whereas, Tbet expression remained at basal levels in the majority of cells activated with low concentrations of pPCC (Fig. S1).

Following activation with a weak TCR-signal (0.1μM pPCC) GATA3 ^Hi^Tbet^Lo^ cells predominated (Fig. 1A). As the strength of pPCC stimuli was increased, the proportion of GATA3^Hi^ cells decreased and proportions of Tbet^Hi^ cells increased, in a signal strength dependent manner, with GATA3^Lo^Tbet^Hi^ cells predominating following stimulation with a strong TCR-signal (10μM pPCC) (Fig 1A). In parallel the patterns of canonical Th1 and Th2 effector cytokines IFNγ and IL-4, closely replicated patterns of Tbet and GATA3 expression (Fig. 1B). At intermediate doses of pPCC, cells exhibited increased levels of both GATA3 and Tbet expression in comparison to naïve cells but did not produce corresponding levels of effector cytokines (Fig. 1A-D). Indicating a potential functional blockade to effector cytokine production or differentiation towards an alternate T-helper lineage. Analysis of Foxp3 and RORγt expression, indicative of differentiation to Treg or Th17 lineages, revealed an absence of RORγt (Fig. S2A). Whereas Foxp3 expression was upregulated in a proportion of activated CD4+ T cells in a manner inversely proportional to the strength of TCR signalling (Fig. S2A).

**Figure 1:**
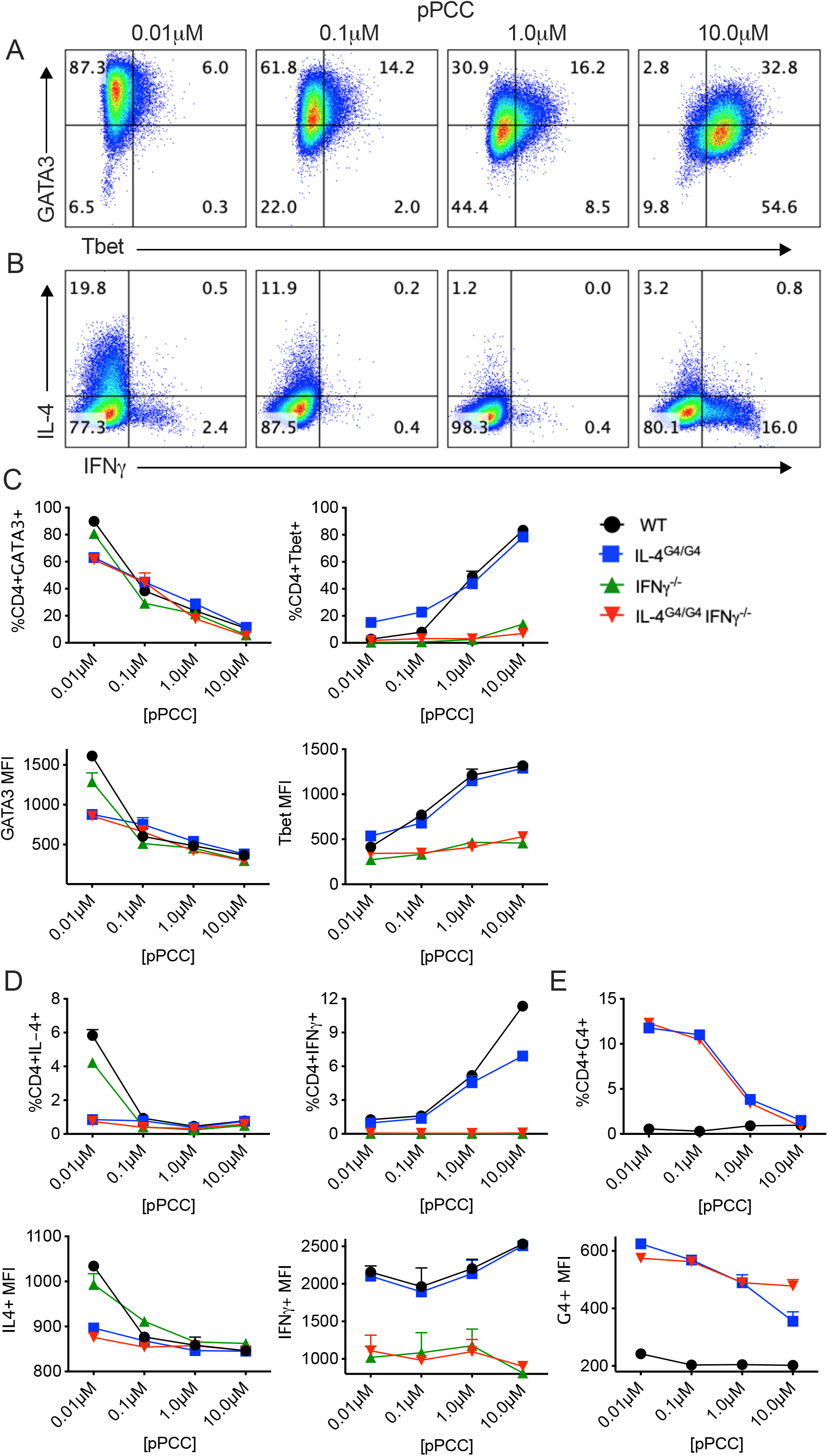
Th1 differentiation requires exogenous signals while Th2 differentiation is induced by an endogenous program. Naïve 5CC7 CD4^+^ T cells were stimulated with P13.9 antigen-presenting cells in the presence of 0.01–10 μM pigeon cytochrome C peptide (pPCC) for 4 days under in vitro conditions. Cells were restimulated with PMA and Ionomycin. (A) Tbet and GATA3 and (B) IL-4 and IFNγ expression was determined following intracellular staining. (C-D) Wildtype (WT), IL-4^G4/G4^, IFNγ^−/−^, IL-4^G4/G4^ IFNγ^−/−^ cells were cultured as in (A). (C) Frequencies and mean fluorescent intensities (MFI) of CD4+ cells expressing Tbet and GATA3 were calculated. (D) Frequencies and MFI of CD4+ cells expressing IL-4, IFNγ, and (E) G4 were calculated. (C-E) Error bars indicate mean ±SEM, n=3; experiments were performed at least three times with consistent results.

### Roles of IL-4 and IFNγ During TCR Signal Strength-Driven Differentiation

To dissect the roles of autocrine IL-4 and IFNγ signalling in TCR-driven differentiation, we analysed the activation of Th-naïve cells from wildtype (WT), IL-4^G4/G4^, IFNγ^−/−^, and IL- 4^G4/G4^IFNγ^−/−^ double-deficient animals. Here, use of the IL-4^G4/G4^ reporter system allows analysis of IL-4 loci expression activity in a functional *IL4* knockout model [43]. Analysis of GATA3 expression indicated that stimulation with a weak TCR-signal led to GATA3^Hi^ expression, even in the absence of IL-4 (Fig. 1C). Again, the proportions of GATA3^Hi^ cells decreased as levels of TCR stimulation were increased (Fig. 1C). Indicating, Th2 differentiation occurred independent from IL-4, but relative to the level of TCR-signalling received. Additional analysis, of GATA3 expression following weak TCR stimulation showed that the proportion of GATA3^Hi^ cells and GATA3 MFI, was closely linked with the ability to produce IL-4, as both WT and IFNγ^−/−^ cells expressed higher levels of GATA3 than IL-4^G4/G4^ or IL-4^G4/G4^ IFNγ^−/−^ cells (Fig. 1C & S3A). Indicating, IL-4 was not required for the induction of GATA3, but autocrine IL-4 production resulted in enhanced GATA3 expression. As in WT cells, no RORγt expression was induced in the absence of IL-4 and IFNγ, and Foxp3 expression was observed to negatively correlate with signal strength in IL-4^G4/G4^ IFNγ^−/−^ cells (Fig. S2). Using G4 expression as a proxy for IL-4 production, we confirmed that, in the absence of IL-4, *IL-4* gene expression was induced in a manner relative to the strength of TCR signal received (Fig. 1E). Further, demonstrating, that in the absence of IL-4, Th2 differentiation was driven by a TCR-signal-dependent, IL-4-independent endogenous program.

In comparison, Th1 differentiation in WT and IL-4^G4/G4^ cells increased following strong TCR-signalling, but was absent in IFNγ^−/−^, and IL-4^G4/G4^IFNγ^−/−^ cells (Fig. 1B & D). Analysis of IFNγ and Tbet expression showed that, as the strength of stimulation increased, the proportion and MFI of cells expressing IFNγ and Tbet increased correspondingly in IFNγ- sufficient populations (Fig. 1C). Indicating that both IFNγ and Tbet were preferentially induced following strong TCR stimulation, which was sufficient for the induction of Th1 differentiation in the presence of autocrine IFNγ production. Hence, under these conditions Th1 differentiation was dependent on both strong TCR signalling and IFNγ , with cells requiring the presence of a signal 3 input to induce differentiation, whereas Th2 differentiation could be induced independent of signal 3 following stimulation with a weak TCR signal.

### Early Transcription Factor Expression is Driven by TCR Signalling

To analyse the early signalling events induced by varying the strength of TCR stimulation, we initially assessed CD69 expression to determine the relative activation status of cells after 24h in culture (Fig. 2A). Expression of CD69 occurred in a digital-like on/off manner relative to the strength of TCR stimuli used during activation (Fig. 2B). As in Allison *et al.*’s [44] analysis of cells in the CD69^+^ ‘on-state’, CD69 MFI displayed a graded analogue response proportional to the TCR signal received during activation, indicative of an ability to respond in a quantitative manner to TCR inputs at early timepoints (Fig. 2C). A comparison of WT and IL-4^G4/G4^IFNγ^−/−^ CD4+ T cells revealed that at 24h post-activation, and prior to the induction of division, an increase in both GATA3 and Tbet was evident (Fig. 2D). Here, GATA3 and Tbet expression was induced relative to the strength of the TCR signal received, with weak TCR signalling inducing increased GATA3 expression, and strong TCR signalling inducing increased Tbet expression. Furthermore, the early increase in TF expression observed at 24h was independent of the presence of either IL-4 or IFNγ (Fig. 2D).

**Figure 2:**
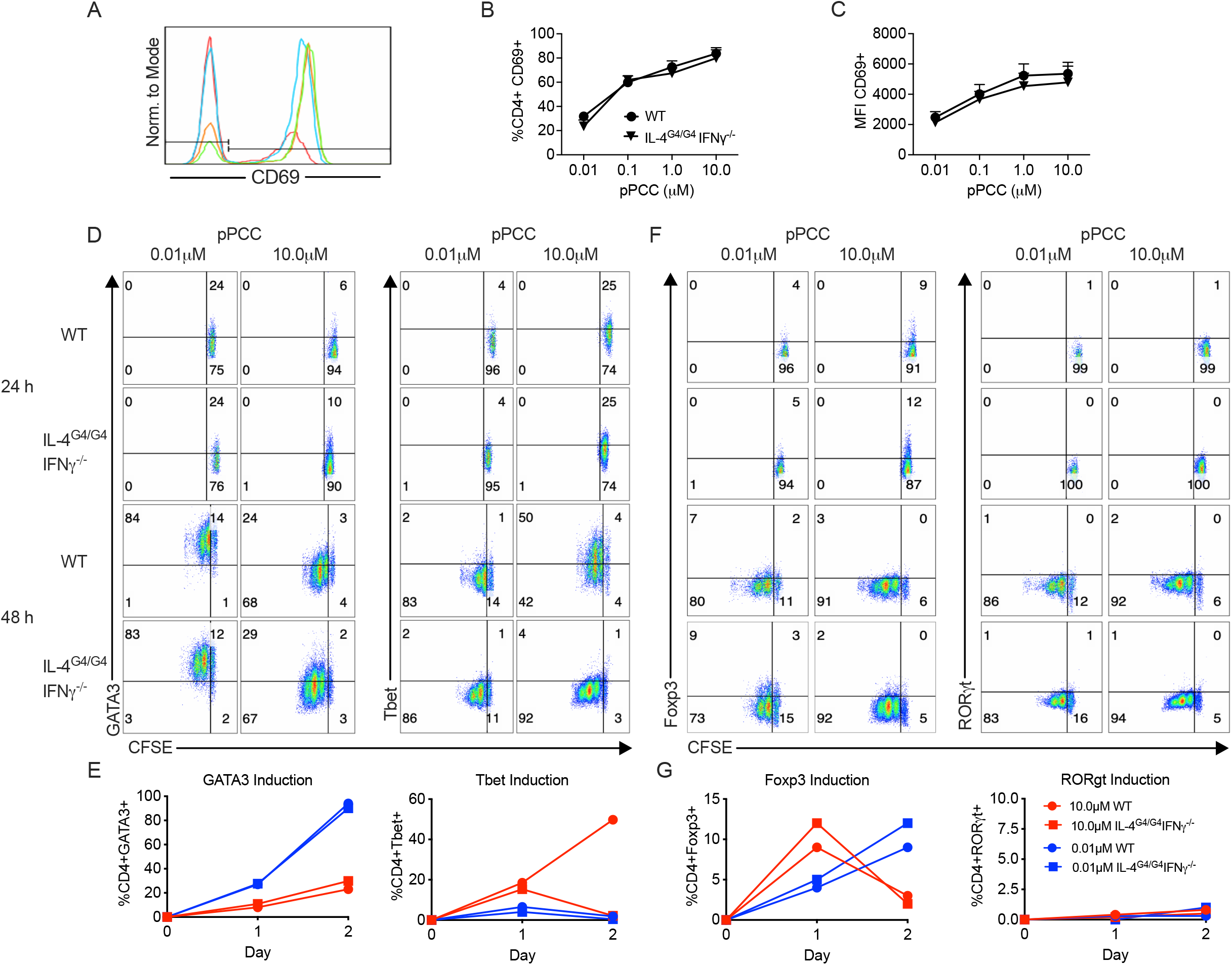
Initial TCR stimulation induces transcription factor expression and cytokine signalling is required for maintenance of Tbet expression. Naive 5CC7 CD4^+^ T cells were labelled with CFSE and stimulated with P13.9 antigen-presenting cells in the presence of pPCC, as previously described. (A) At 24-h post-stimulation, activation of CD4+ T cells was assessed by flow cytometric measurement of CD69 expression (pPCC: 10.0 μM, green; 1.0 μM, orange; 0.1 μM blue; 0.01 μM, red) and (B) the proportion of cells expressing CD69 and the MFI of expression determined. Induction of transcription factor expression in activated CD4+ T cells and cellular division was assessed by comparing CFSE with GATA3, Tbet, Foxp3, and RORγt. (D) Transcription factor expression by activated CD4+ T cells was quantified over time (h). Experiments were performed twice with consistent results.

At 48h post-activation, levels of GATA3 increased in all conditions, with substantially greater proportions of cells expressing GATA3 following weak TCR stimulation. Tbet expression returned to baseline in cells activated with weak TCR-signals and in the absence of IFNγ (Fig. 2D&E). Similar transient increases in the production of early Tbet have also been noted in previous studies [45; 46; 47]. Together, these results indicate that, the presence of IFNγ during an initial deterministic (0-48h) phase of differentiation was necessary to stabilize and reinforce the early cytokine independent expression of Tbet.

The early kinetics of RORγt and Foxp3 induction were next assessed to determine whether a similar proximal TCR-driven induction occurred (Fig. 2F&G). At 24h post-activation, increased Foxp3 expression was detected and at 48h post-activation with a strong TCR-signal, Foxp3 expression declined. In contrast, the proportion of cells expressing Foxp3 when stimulated with a weak TCR-signal increased at 48h (Fig. 2F&G). RORγt expression was not detected at either time point (Fig. 2F&G), indicating that the upregulation of RORγt was not directly induced by TCR stimulation, in contrast to our observations for Foxp3, GATA3, and Tbet.

### Exogenous IL-12 Enhances Th1 Differentiation Synergistically with TCR Signalling While Retaining Memory of Initial Stimulus

To more closely model *in vivo* conditions where Th1 differentiation generally occurs following activation in the context of a DC providing a Th1 inducing signal 3 cytokine [14]. We simulated the effects of IL-12 acting in a paracrine manner to modulate TCR dependent differentiation. (Fig. 1). A fine-grained analysis of TCR signalling in combination with exogenous IL-12, was conducted by titrating the levels of both pPCC and IL-12 during activation (Fig. 3A & B). The inclusion of IL-12 at higher concentrations led to a striking increase in Th1 differentiation at all levels of TCR stimulation. As the concentration of IL-12 was reduced, the impact of varying the strength of TCR signalling began to become more predominant. At 0.01ng of IL-12, TCR-signal-driven differentiation became the major determinant driving differentiation, with weak TCR-stimulation inducing similar proportions of IL-4^+^ cells to those cultured in the absence of IL-12. Whereas 0.01ng IL-12 in combination with strong TCR-stimulation substantially enhanced IFNγ production and IL-12 contributed in a synergistic manner to Th1 differentiation (Fig. 3A). Further analysis, of IL-4 production showed that regardless of IL-12 levels present, when initially stimulated with a weak TCR-signal, a proportion of cells producing IL-4 was consistently observed (Fig. 3A). Analysis of TF expression additionally revealed the distinct effects of TCR signal strength on GATA3 and Tbet expression independent of IL-12, under all conditions tested (Fig. 3B). Even in the presence of 10ng/ml IL-12, the pattern of GATA3 expression was higher following activation with weak TCR-stimuli and Tbet expression higher following strong TCR-stimuli. Further, the relative ratio of GATA3:Tbet expression was influenced by both the levels of exogenous IL- 12 and the concentration of peptide present, with the strength of the TCR stimulation observed to imprint a lasting transcriptional signature despite the presence of supraphysiological levels of IL-12 (Fig. 3B).

**Figure 3:**
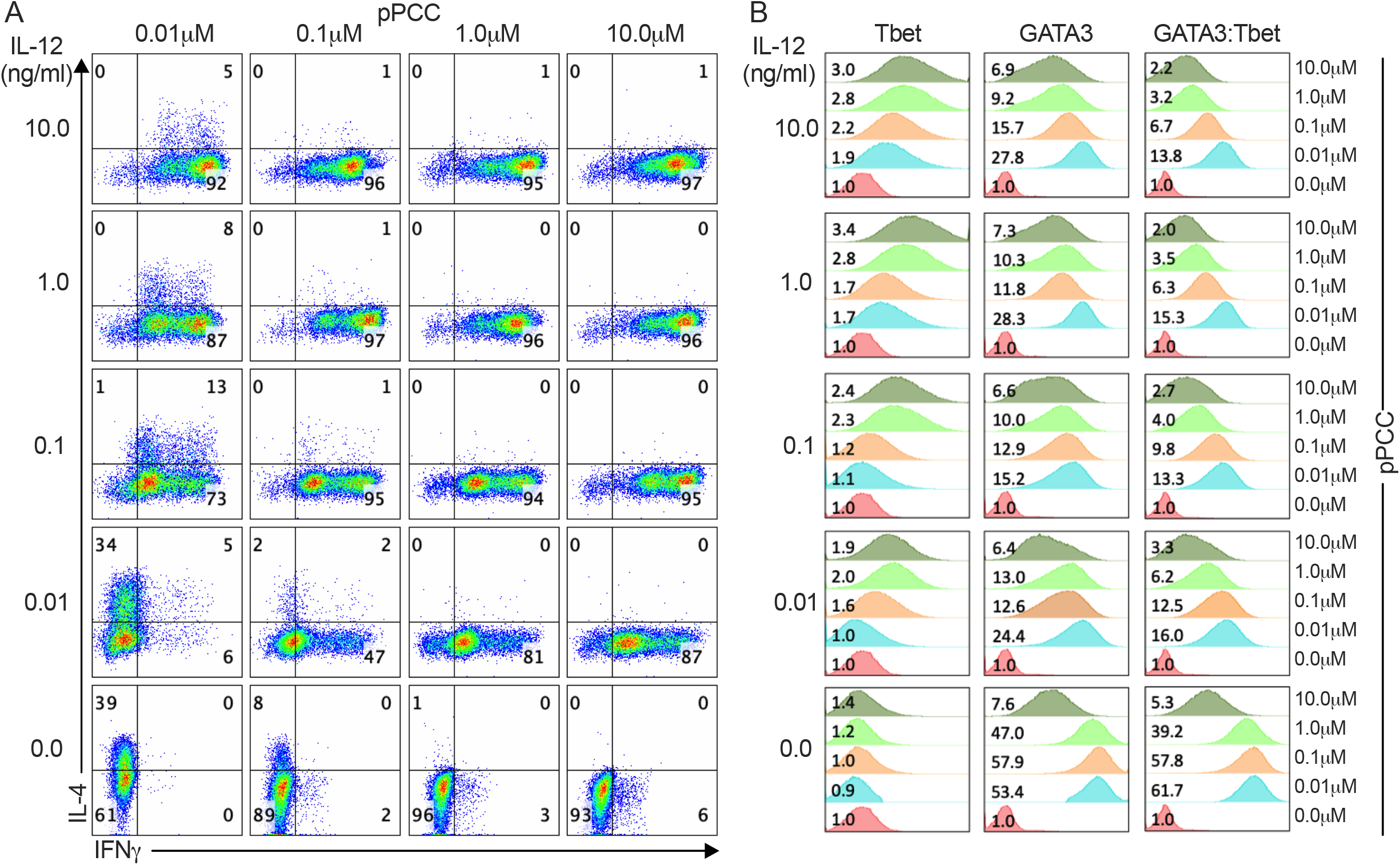
Exogenous IL-12 enhances CD4+ Th1 cell differentiation and synergizes with strong TCR signalling. Naive 5CC7 CD4^+^ T cells were stimulated with antigen-presenting cells in the presence of 0.01–10.0 μM pPCC for 4 days under in vitro conditions, and cells were then re-stimulated. WT CD4+ T cells were cultured in the presence of IL-12, as indicated, and stained for IL-4 and IFNγ expression (A) and Tbet and GATA3 expression (B); inset values represent fold change in MFI compared to unstimulated cells (0.0 μM). Experiments were performed three times with consistent results.

### Initial TCR Signalling Has Lasting Impacts on Cellular Differentiation Profiles

To further examine whether initial TCR signalling imprinted a lasting memory on Th-naive cells, we next analysed the impact of exogenous IL-4 on differentiation. Th-naive cells from WT and IL-4^G4/G4^IFNγ^−/−^ were activated in the presence of a supraphysiological concentration of IL-4 (100U/ml). In the presence of IL-4, a significant increase in the proportion of GATA3^Hi^ cells was observed in IL-4 deficient populations (Fig. 4A,B,S4A) and GATA3 expression was comparable to that induced in WT cells without exogenous IL-4 following activation with a weak TCR-stimuli (Fig. S4A). Analysis of WT cells activated with a weak TCR-stimuli in the presence of IL-4 revealed no significant difference in either the proportion of GATA3^Hi^ cells or the MFI of GATA3 in comparison to WT cells cultured alone (Fig. S4A), potentially indicating an upper limit to GATA3 expression levels. In comparison, IL-4^G4/G4^IFNγ^−/−^ cells activated with a weak TCR-stimuli exhibited decreased levels of GATA3 expression compared to WT cells, which was recovered by the addition of exogenous IL-4. The TCR-signal-strength-dependent effects on Tbet expression remained evident, as Tbet expression continued to be positively associated with the initial strength of TCR signal encountered (Fig. 4C). In both WT and IL-4^G4/G4^IFNγ^−/−^ cells, IL-4 (Fig. 4D) and G4 (Fig. 4F) expression was enhanced by the presence of exogenous IL-4 during differentiation. In the absence of exogenous IL-4, IFNγ-sufficient WT cells exhibited a TCR-dose-dependent decrease in IL-4 production, with a corresponding rise in Tbet expression (Fig. 4C). Whereas, in the absence of IFNγ, G4 expression did not decrease in IL-4^G4/G4^IFNγ^−/−^ cells stimulated with a strong TCR- stimuli (Fig. 4F). IFNγ production was only detectable in WT cells cultured without the addition of IL-4 (Fig. 4E). Tbet expression was more closely correlated with the strength of stimulation received, as expression decreased in the presence of both endogenous and exogenous IL-4 and in the absence of IFNγ in a TCR-dose-dependent manner (Fig. 4A&C). Here, the absence of IFNγ in combination with exogenous IL-4 was found to most significantly repress Tbet expression (Fig. 4C&S4B)

**Figure 4.**
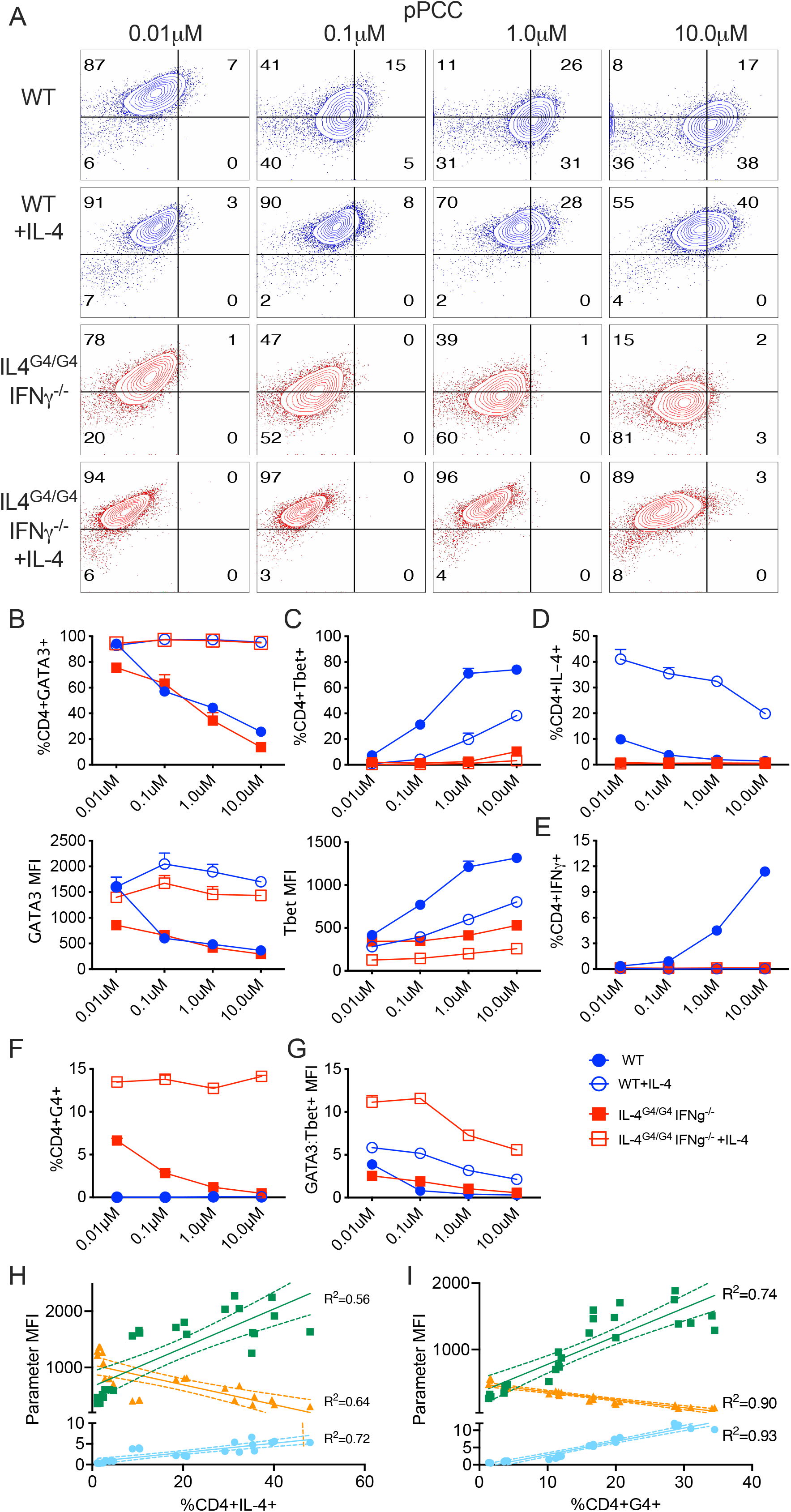
**Exogenous cytokines enhance Th2 CD4+ T cell differentiation, following integration of TCR signalling.** (A) Naive WT and IL-4^G4/G4^ IFNγ^-/-^ 5CC7 CD4+ T cells were cultured with APC in the presence of pPCC as previously described, exogenous IL-4 was added to WT and IL-4^G4/ G4^ IFNγ^-/-^ cells, as indicated. At day 4 cells were then restimulated and Tbet and GATA3 expression analysed. Proportions and MFI of (B) GATA3 and (C) Tbet expressing cells were determined and the proportion of cells and expressing (D) IL-4, (E) IFNγ and (F) G4 was determined. (G) The Ratio of GATA3 MFI to Tbet MFI was calculated on a per cell basis. GATA3 MFI (▪), Tbet MFI (▲) and GATA3:Tbet MFI (●) levels were compared with %IL-4 expression for (H) WT cells, or to G4 expression for (I) IL-4^G4/G4^ IFNγ^-/-^ cells and linear regression analysis conducted. (A) Data shown is representative of two separate experiments. (B-G) Data shown is representative of n=3; error bars indicate mean ± SEM. Experiments were performed twice with consistent results. (H,I) Simple linear regression analysis was conducted and R^2^ values determined as indicated; dashed lines show 95% confidence intervals.

In response to the activation of WT cells in the presence of exogenous IL-4, the TCR- dose-dependent induction of Tbet remained. While the levels of Tbet expression were significantly reduced, the presence of exogenous IL-4 largely exerted its effect by upregulating the expression of GATA3, as opposed to inhibiting Tbet expression (Fig. 4A & B). With the help of linear modelling, we evaluated the relative production of IL-4 due to the presence of either GATA3 or Tbet and tested two model scenarios (Fig. S5). In both WT and IL-4^G4/G4^FNγ^−/−^ cells, the presence of Tbet showed a non-significant negative association with IL-4 production. However, upon the addition of IL-4, the coefficient increased and became statistically significant. As the model fitted evaluated both the effect of GATA3 and Tbet simultaneously for each condition, we observed a stronger negative association with Tbet compared to the positive association with GATA3 in the presence of exogenous IL-4. In WT and IL-4^G4/G4^IFNγ^−/−^ cells, GATA3 had a stronger positive association with IL-4 production than with Tbet, indicating the need for a model which incorporates both GATA3 and Tbet to predict the probability of cytokine production levels following TCR-dose-dependent differentiation. Our findings show that the co-expression of high levels of GATA3 and Tbet efficiently inhibited IFNγ production and were more permissive for IL-4 production (Fig. 4A,D-F). Together, these results indicate that, whilst GATA3 is required for Th2 differentiation [48; 49], the induction of additional factors, including the co-expression of Tbet, plays a significant role in differentiation and determining the ability of an individual cell to produce IL-4.

To better model the comparative influence of GATA3 and Tbet on differentiation, the ratio of GATA3 to Tbet was calculated on a per cell basis (Fig. 4G). A comparison of GATA3:Tbet with respect to TCR stimuli revealed a dose-dependent effect of TCR signalling on TF expression, indicating that the relative GATA3:Tbet expression level may control effector cytokine production. To test this hypothesis further, we combined IL-4 or G4 expression data from WT or IL-4^G4/G4^IFNγ^−/−^ cells respectively, using cells stimulated both with and without exogenous IL-4 at all peptide doses, a dataset was generated with a range of expression values for either IL-4 or G4. A simple linear regression analysis comparing GATA3, Tbet, and GATA3:Tbet expression values with regards to IL-4 or G4 expression levels was then performed (Fig. 4H&I) and in both cases, use of the GATA3:Tbet parameter most closely matched the plotted data.

### Analysis of Differential Gene Expression Networks Induced by Modulating TCR Signal Strength

The transcriptomes of CD4+ T cells activated with pPCC over 4 days were assessed without restimulation to preserve the steady-state patterns of gene expression. Analysis of signature Th-cell-associated genes revealed a graded response between Th2 and Th1 differentiated cells, in which TFs, cytokines, cytokine receptors, and lineage-associated surface receptors largely matched those of previously reported Th molecular phenotypes [50](Fig. 5A).

**Figure 5:**
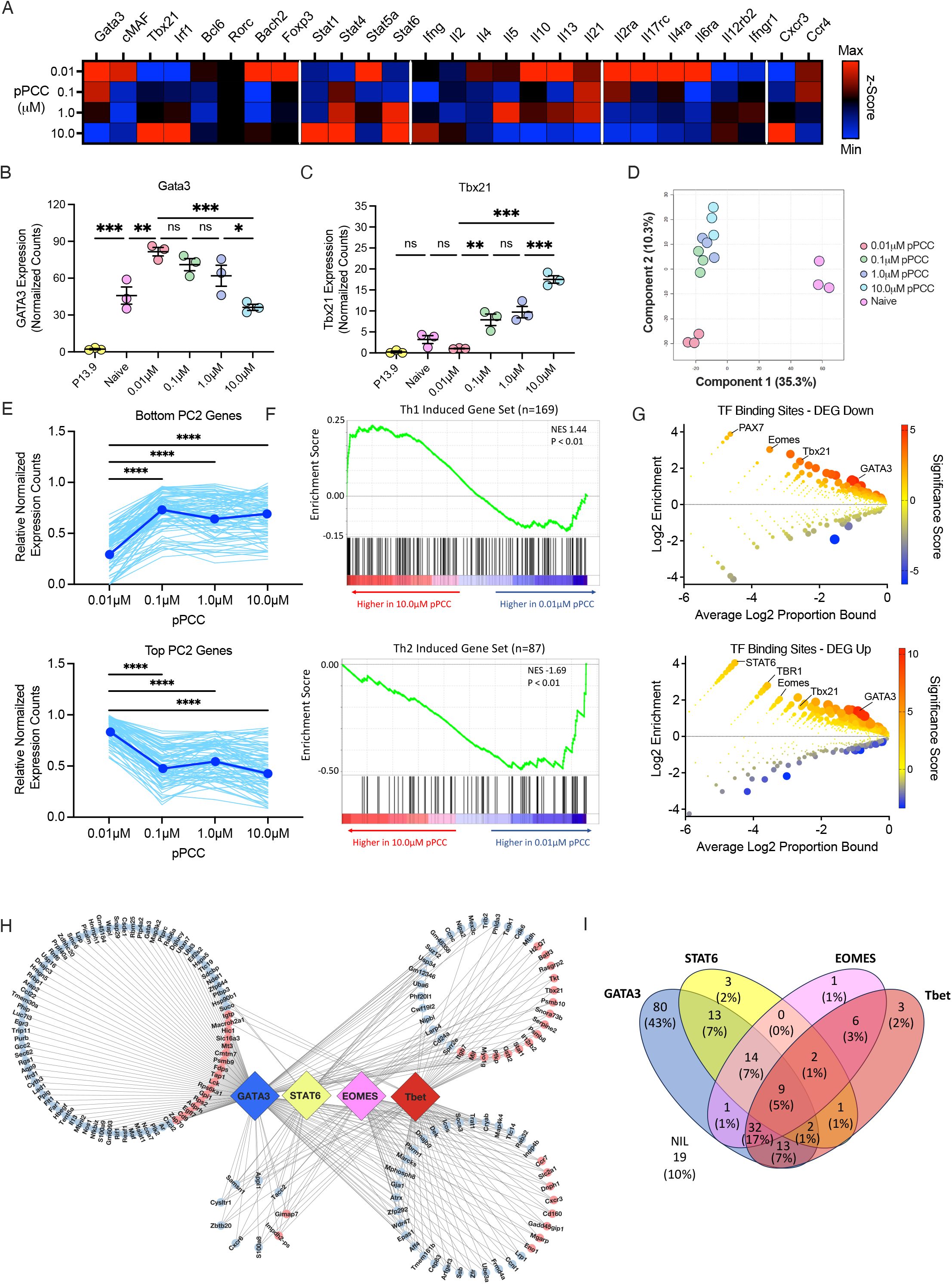
**GATA3 and Tbx21 are central regulatory factors in a differential gene expression network** Naïve CD4^+^ T cells were stimulated with 0.01 μM-10.0 μM pPCC for 4 days under in vitro conditions and cells were processed for transcriptome analysis by RNAseq without restimulation. (A) Comparative expression levels of selected Th-associated markers of differentiation. Comparison of (B) Gata3 and (C) Tbx21 gene expression normalized to read counts per million transcripts (CPM). (D) Partial least squares discriminant analysis of Th-naïve and stimulated CD4+ T cells . (E) Normalized expression values for bottom and top 1% of genes contributing to PC2 in PLSDA plot (D) were compared to TCR signal strength (light blue) and mean value plotted (dark blue). (F) GSEA showing enrichment of Th1-induced genes (top) and Th2-induced genes in comparison of transcriptomes of CD4+ T cells. (G) Enrichment analysis of transcription factor (TF)-binding sites associated with downregulated (top) and upregulated (bottom) genes in 10.0μM vs. 0.01μM pPCC groups. (H) Network diagram showing GATA3, STAT6, EOMES, and Tbet binding sites of DEGs. Edges indicate the presence of a TF-binding site, blue nodes indicate downregulated genes, red nodes indicate upregulated genes. (I) Proportions of DEGs with binding sites for the TFs profiled. (B&C) Error bars indicate mean ±SEM, n = 3. Statistical analysis was performed by 1-way ANOVA with Tukey multiple comparison testing. * = p < 0.05, ** = p < 0.01, *** = p < 0.001, ns = nonsignificant.

Patterns of *Gata3* and *Tbet* expression were significantly differentially regulated at the transcriptomic level and were preferentially induced by weak versus strong signalling, respectively (Fig. 5B&C). The transcriptomes from pPCC-stimulated cells were compared with those of Th-naive cells a PL-SDA model was used to determine the primary component features present (Fig. 5D). Component one effectively segregated the Th-naïve cells from the activated CD4+ T cells, whereas component two accurately discriminated between the activated CD4+ T cell populations, revealing a distribution closely correlating with TCR signal input. Here, we extracted the most positive and negative 1% of genes from component 2 to determine which genes most significantly contributed to the expression axis (Fig. 5E). The negative gene set increased in a graded manner concomitant with TCR signal strength and consisted of many Th1 associated genes including *Tbx21, Il12rb* and *Cxcr3.* In contrast, the positive gene set showed an opposite pattern of behaviour, but was not found to include those genes commonly associated with Th2 differentiation. Gene ontology analysis revealed several enriched pathways for the negative gene set including pathways associated with T cell activation and type-II-interferon responses, whereas the positive gene set showed enrichment for only the positive regulation of organelle organization (Fig. S6A). To further characterise transcriptomic differences, we utilised gene set enrichment analysis (GSEA) which showed a significant enrichment in Th1 signature genes following activation with μM pPCC, whereas a significant enrichment in Th2 genes was associated with 0.01μM pPCC stimulation (Fig. 5F), further confirming the previously observed protein signatures.

To assess the overall impact of the differentially expressed TFs on the programming of differentiation, those TFs present amongst all DEGs were identified (Fig. S6B-D), and associations analysed based on the patterns of expression. Unbiased clustering indicated the presence of two important Th-associated modules (Fig. S6B), consisting of a Th1 module (*Tbx21, Irf1, Stat1, Batf3, Hic1, Eno1,* and *Zfpm1*) and a Th2 module (*Gata3, Maf, Epas1, Aff4, Suz12,* and *Zfp292*). These differentially expressed genes showed a consistent pattern of either positive or negative correlation with TCR signal input. A STRING network was constructed from the differential TFs to assess the interactions between the TF nodes, which further highlighted the close relationship between key regulatory proteins (Fig. S6C). To better assess the potential for intergenic regulatory effects by key T-helper associated TFs, transcription-factor-binding sites (TFBS) were identified in the regulatory regions of the DEGs. In those genes associated a Th1 signature (Fig. 5G), we found a significant enrichment in the binding sites for the Th-associated TFs Tbet, Eomes, and GATA3. In comparison, in the DEGs associated with Th2 differentiation (Fig. 5G), the TFBS for GATA3 and STAT6 were significantly enriched and those for Tbet and EOMES were also present, but at lower levels. Gene–TF interactions were visualized with a network diagram (Fig. 5H), and the number and proportion of putative binding sites were determined (Fig. 5I). We determined GATA3 had the potential to regulate the largest number of DEGs, with a substantial number of GATA3 binding sites present in both Th1-and Th2-associated genes. Whilst a large number of potential Tbet binding sites were present amongst the DEGs, comparatively few genes were predicted to be regulated by Tbet alone. Together, the DEG expression and TFBS network data suggest a model in which increasing TCR signal strength induces changes in Th differentiation patterns through modulating both the number of genes expressed and the level of gene expression in a complex network of coregulatory TF binding.

### Ratiometric Expression of GATA3 and Tbet Quantitatively Governs Cytokine Expression in Differentiated Th1 and Th2 Cells

To better understand the relationship between GATA3 and Tbet expression levels and their quantitative effects on the ability of differentiated CD4+ T cells to produce IFNγ or IL-4. We reverted to using a model system unbiased by the addition of exogenous cytokines. Here, effector cytokine production was compared to expression of GATA3, Tbet and to GATA3:Tbet levels present in individual cells (Fig. 6A). As expected, IFNγ production corresponded with higher levels of Tbet expression, and IL-4 production corresponded with increased GATA3 expression. Furthermore, use of GATA3:Tbet as a parameter to assess the functional relationship between TFs and cytokine production, revealed high and low levels of GATA3:Tbet closely corresponded with the production of IL-4 or IFNγ respectively (Fig. 6A). The quantitative relationships between GATA3, Tbet, and GATA3:Tbet levels with IFNγ or IL-4 production in cytokine expressing cells was then assessed. Subpopulations of cells initially activated with 10.0μM or 0.01μM pPCC were gated based on their production of IFNγ (IFNγ^Hi^/IFNγ^Lo^) or IL-4 (IL-4^Hi^/IL-4^Lo^) (Fig. 6B). Following activation with the Th1- polarizing 10.0μM pPCC, levels of Tbet were found to be elevated in IFNγ producers, with increased Tbet expression observed consistent with the degree of IFNγ production (Fig. 6C); whereas GATA3 was only significantly increased in IL-4^Hi^ producers in comparison to non-producers (Fig. 6C). Comparison of GATA3:Tbet to cytokine production increased the significance of the associations between TF levels and cytokine production, with a significant and consistent trend from IFNγ^Lo^ to IL-4^Hi^ groups observed. Activation with 0.01μM pPCC also revealed a significant quantitative relationship between IFNγ production and the levels of Tbet expression and IL-4 production with GATA3 expression. Again, the use of the GATA3:Tbet parameter as a predictor of a specific cell’s ability to produce either IFNγ or IL-4 yielded a closer association, with significant differences in GATA3:Tbet detected for each of the populations analysed (Fig. 6C).

**Figure 6:**
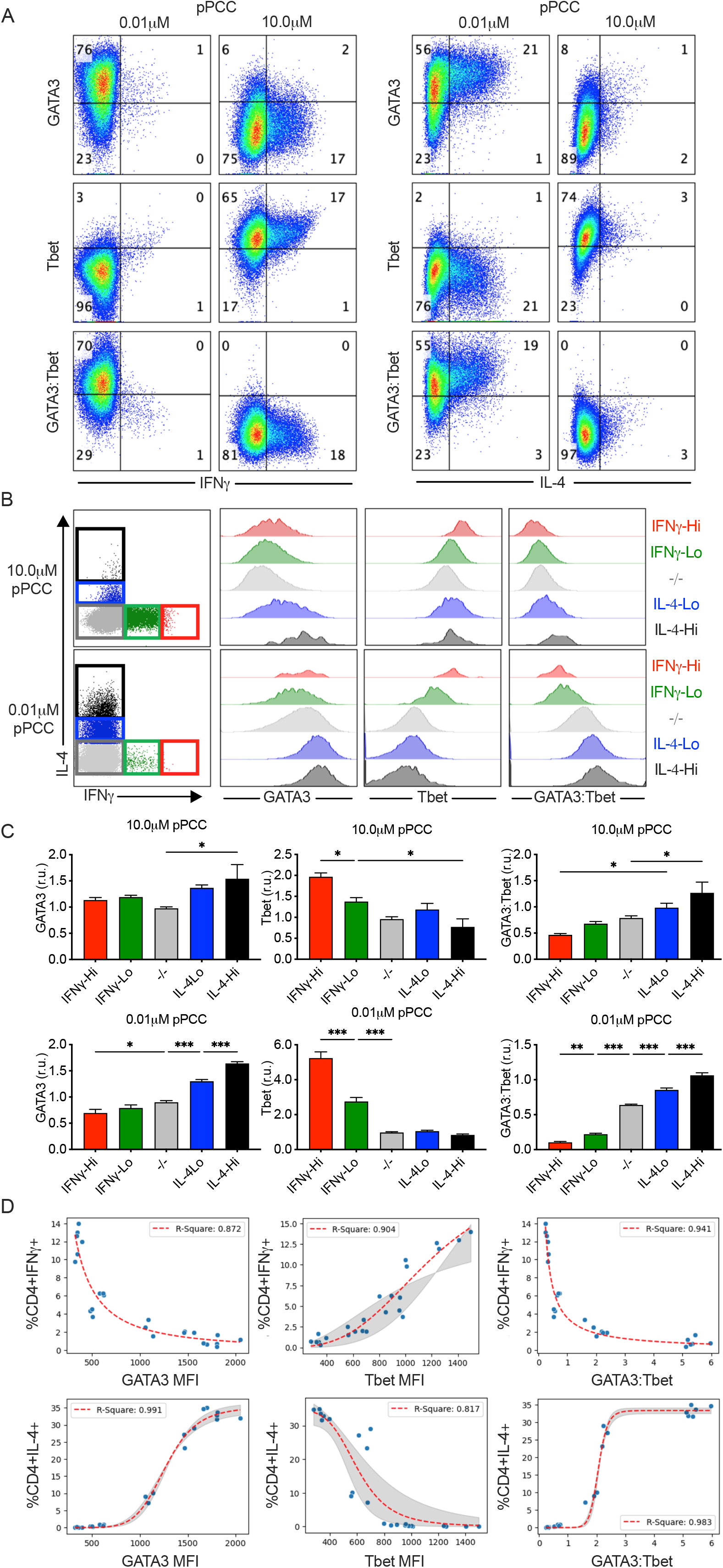
**Cytokine expression in effectors cells is related to levels of transcription factor expression** Naïve 5CC7 CD4^+^ T cells were stimulated with P13.9 antigen-presenting cells in the presence of 0.01 μM or 10.0 μM pPCC for 4 days under *in vitro* conditions. Following restimulation, the levels of GATA3, Tbet, and GATA3:Tbet expression ratio were compared to (A) IFNγ expression or IL-4 expression. (B) CD4+ T cells were gated based on the expression of IL-4^Hi^ (black), IL4^Lo^ (blue), IFNγ (red), IFNγ (green), or non-expressors (grey). Comparative expression of GATA3, Tbet, and GATA3:Tbet was plotted for each population. (C) Expression levels relative to total population of GATA3, Tbet, and GATA3:Tbet expression were calculated in relative units (r.u.) and compared for each cytokine-expressing population. (D) Correlation of IFNγ+ or IL-4+ frequency with GATA3, Tbet, or GATA3:Tbet expression in 5CC7 stimulated with 0.01 μM, 0.1 μM, 1.0 μM, or 10.0 μ M pPCC (A-D). Data are representative of three experiments and are shown as means ±SEM (C&D), n = 6. Statistical analysis was performed by 1-way ANOVA with Tukey multiple comparison testing. Red line in (D) shows the best fit to the data obtained via model fitting conducted using Hill’s equation. The shaded region indicates the 95% confidence interval.

To further characterize the quantitative relationship between TF levels and cytokine production, we gated cells activated with either strong- or weak-TCR-stimuli into three populations (Hi, Int, or Low) based on expression levels of GATA3, Tbet, or GATA3:Tbet (Fig. S7A). The relative production of IL-4 showed positive associations with GATA3 and GATA3:Tbet and a negative association with Tbet expression (Fig. S7B). Here, the significance of the associations was stronger for those cells activated with weak TCR-stimuli (Fig. 6B). In comparison, the relative production of IFNγ compared to TF expression showed a reversal of the trend observed for IL-4. Except in the case of GATA3-Hi and GATA3-Int groups, as following activation with a strong TCR-stimuli, IFNγ production was positively associated with GATA3 (Fig.S7C). Heatmapping of IL-4 or IFNγ production levels onto GATA3 versus Tbet expression plots confirmed that those cells producing the highest levels of IFNγ also expressed greater amounts of both GATA3 and Tbet (Fig. S7D. Additionally, analysis of the normalized proportions of IL-4^+^ and IFNγ^+^ cells present in comparison to GATA3, Tbet, and GATA3:Tbet levels largely replicated these associations, indicating that the relative expression of GATA3:Tbet had the strongest associations with cytokine production, in terms of probability of expression (Fig. S7E&F) and levels expressed (Fig. S7B&C).

To ascertain whether the expression levels of GATA3 and Tbet have a causal role in the ability of cells to produce cytokines at the population level, we extended our analysis to include those cells activated with intermediate levels of TCR stimulation. These cells had previously been shown to exhibit increased TF expression in comparison to naïve cells but failed to produce the corresponding levels of effector cytokines. We initially employed an empirical dose-response curve to model the relationship between TF expression and the proportion of cells within each population that produced cytokines upon restimulation (Fig. S8), as in Helmstetter *et al.* [42]. Analysis of associations between GATA3, Tbet, and GATA3:Tbet with IFNγ production, revealed that GATA3:Tbet provided a closer association. In contrast, when we fitted for IL-4 production, Tbet had the weakest association, and GATA3:Tbet again had the strongest fit for the data. To better model the sigmoidal appearance of the data, we utilized a curve fitting model incorporating the Hill equation (Fig. 6D). Again, we saw that, for IFNγ production the GATA3:Tbet function resulted in the greatest R^2^ value, followed by Tbet and GATA3. For IL-4 production, GATA3:Tbet and GATA3 allowed us to model the data very closely, with R^2^ values of 0.991 and 0.983, respectively.

Taken together, our findings show a strong causal relationship between both GATA3 and Tbet levels in predicting the ability of an individual cell to produce either IL-4 or IFNγ and demonstrate that the application of a GATA3:Tbet function to quantitate the relative expression of TF’s provides a more accurate modality for determining the probabilistic frequency of effector cytokine production.

*Duration of TCR-mediated signalling as a mechanism for modulating Th1/Th2 differentiation* As previous *in vivo* studies have demonstrated that the strength of the TCR signal encountered corresponds with the average duration of interactions between T cells and APCs [28; 51]. Here, we aimed to test the importance of TCR signal duration in determining the outcome of differentiation. We initially compared the relationship between the strength of TCR signalling with the duration of interactions during the first 2 h of culture. Th-naive cells interacting with APCs under high-dose conditions were observed to have significantly greater bursts of Ca^2+^ flux upon encountering an APC (Fig. 7A & B; SMovie1) and increases in interaction times compared to cells activated under low-dose conditions (Fig. 7C). To effectively modulate the duration of TCR signalling, Th-naive cells were activated with pPCC loaded APC’s, and an anti-MHCII blocking antibody was subsequently added to disrupt TCR:pMHCII interactions [52]. Cells were then cultured until 4 days post-activation, and T-helper differentiation and function determined (Fig. 7D&E). As previously observed, in the absence of MHCII blocking, signal-strength-dependent differentiation into Th1 and Th2 subsets was evident. Intriguingly, stimulation for only 2h induced a similar pattern of signal-strength-dependent differentiation, with high-dose stimulation inducing Th1 differentiation and low-dose stimulation leading to Th2 polarization (Fig. 7D). These results were confirmed by analysis of Tbet and GATA3 expression (Fig. 7E). Additionally, cells activated for 12h and 24h also showed similar patterns of dose-dependent differentiation into Th1 and Th2 effector lineages (Fig. 7D&E). Entry into cellular division was also assessed, and following 2h of stimulation, >90% of all surviving cells had not divided (Fig. 7F). Demonstrating that entry into division was not required for the establishment of a distinct pattern of lineage-defining master regulators of differentiation or for the ability of CD4+ T cells to produce the canonical cytokines associated with effector differentiation upon recall stimulation. Analysis of cell numbers following differentiation revealed that cellular expansion and survival was closely linked to both the strength of stimulation received and the length of time cells received stimulation (Fig. S9A&B). When the duration of TCR stimulation was reduced, cells progressed through fewer rounds of division, and the overall rates of cell survival post-differentiation were decreased in a manner directly related to the initial length of interaction (Fig. 7F&S9B). Taken together, our data suggest that the strength of TCR stimuli and the initial duration of TCR stimulation represent two distinct signalling inputs, where the strength of the initial TCR stimulus initiates a program of differentiation at a very early time point, independent from the requirement for cell division. Whereas ongoing stimulation through the TCR is required to drive CD4+ T cells into division, and additionally regulates the number of rounds of expansion cells progress through and modulates cell survival rates.

**Figure 7:**
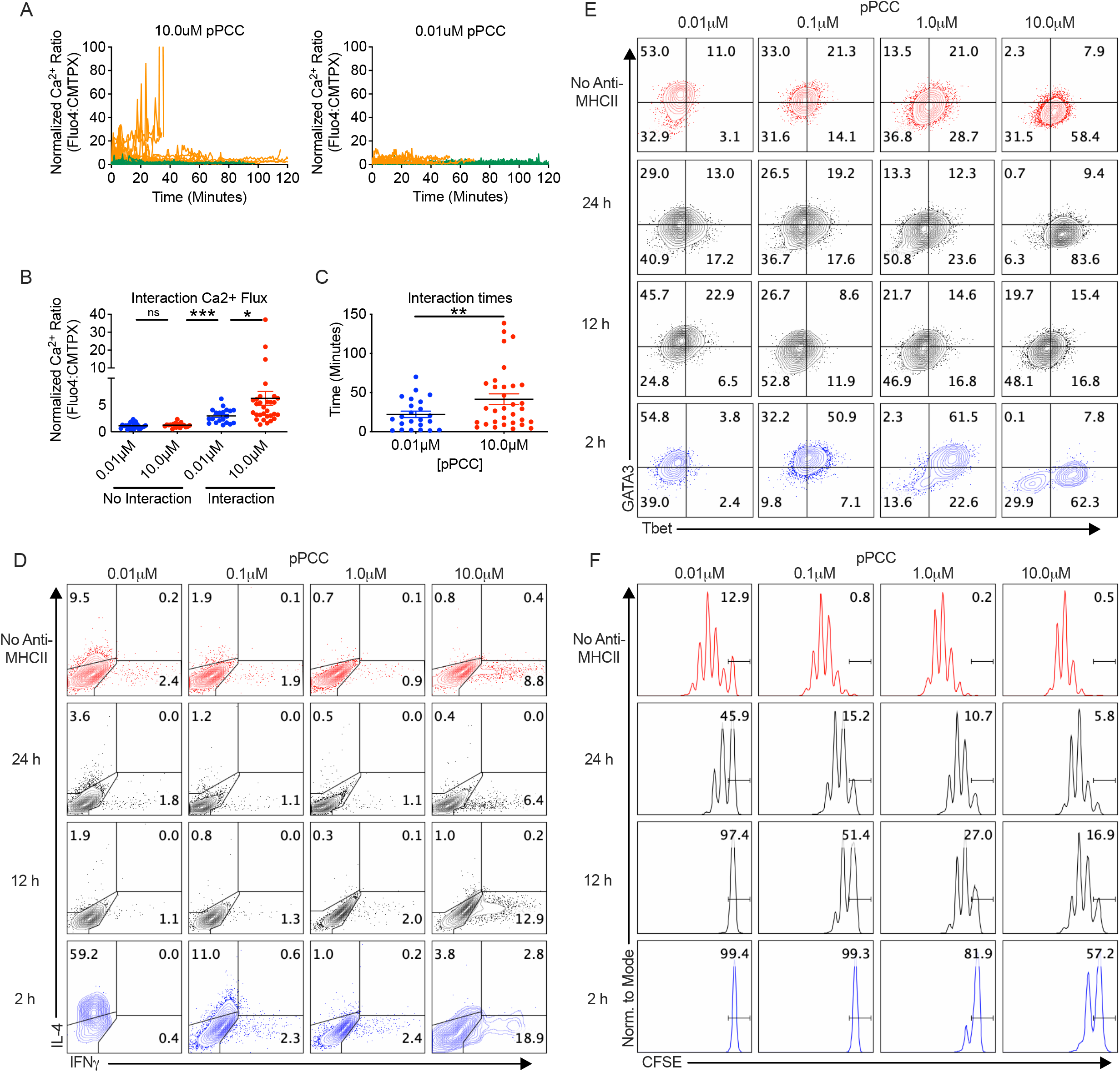
**TCR signal strength controls differentiation while TCR signal duration controls division** Fluorescently labelled naïve 5CC7 CD4^+^ T cells were stimulated with P13.9 antigen-presenting cells (APCs) in the presence of either 0.01 μM or 10.0 μM pPCC for 2 h under in vitro conditions. (A) Individual 5CC7 Ca^2+^ flux tracings for cells interacting with APC’s are shown for duration of T cell:APC contact (red) and non-interacting (blue. (B) Normalized mean Ca2+ flux values for each interaction. (C) Individual 5CC7 cell interaction times with APC’s. (D-F) CFSE labelled CD4+ T cells were activated with APC’s and the duration of TCR stimulation was controlled by the addition of anti-MHCII antibodies at 2h to halt interactions. Differentiation was determined at day 4 post-activation via analysis of (D) IFNγ and IL-4; or (E) Tbet and GATA3. (F) Induction of cell division was assessed on day 4 post-activation by CFSE dilution analysis. All experiments were performed at least twice with consistent results. Statistical analysis was performed via 1-way ANOVA with Tukey multiple comparison testing (B), or Students t-test (C). * = p < 0.05, ** = p < 0.01, *** = p < 0.001, ns = non-significant.

## Discussion

Traditional models of differentiation are based on a reciprocal system of regulation controlled by the qualitative effects of polarizing cytokines which drive the induction of lineage-defining master TFs. Whilst, in turn, downregulating opposing signalling pathways and repressing the expression of those TFs responsible for driving alternate T-helper fates [49; 53]. Recently, data supporting a more nuanced model of differentiation has emerged, in which CD4+ T cell polarization constitutes a continuum of states which display a greater degree of plasticity than initially supposed [54; 55; 56; 57]. Activated human and murine CD4+ T cells have been shown to exhibit the ability to co-express varying levels of regulatory TFs including GATA3 and Tbet, under a variety of conditions [50; 58; 59; 60]. Furthermore, co-expression of the Treg defining TF Foxp3 with Tbet or GATA3 has been shown to help mediate specific regulatory responses [61; 62]. Here, we put forward a model of differentiation whereby initial stimulatory signals imparted by APCs both activate Th-naïve cells and imprint instructions for early TF expression through the quantitative integration of TCR associated signals. Subsequently the presence of autocrine and paracrine cytokine signalling acts downstream, to either enhance or repress early TF expression to further tune the outcome of differentiation. Our data suggest that following activation the process of differentiation results in the induction of a continuum of T-helper states based on the relative expression of lineage-defining TFs, which in turn determines the probabilistic frequency of effector cytokine production on a per cell basis.

IL-4 is largely thought of as being the key driver and the third signal required in the activation/differentiation cascade that results in Th2 differentiation. Here, we establish that in the absence of IL-4, Th2 differentiation can be efficiently induced following activation with a weak TCR-stimuli. As in the absence of IL-4, both the upregulation of GATA3 and induction IL-4 gene expression through use of the G4 reporter were observed. Previous findings showing that IL-4-independent Th2 differentiation occurs under *in vivo* conditions [6; 7; 8; 9; 10], combined with the constitutive expression of GATA3 at low levels in Th-naïve cells, has given rise to the hypothesis that Th2 differentiation may be the result of a default differentiation pathway [14]. However, data presented here indicates that the IL-4 independent ability of Th-naive cells to upregulate GATA3 expression and differentiate into Th2 cells is the result of a controlled process, regulated by the intensity of TCR associated signalling received during activation. Rather than via a default pathway which broadly induces the upregulation of GATA3 expression in the absence of a repressive cytokine signalling program. Here, we observed CD4+ T cells activated with a strong TCR signal in the absence of polarizing cytokine signalling (IL-4^G4/G4^IFNγ^−/−^) did not default to a Th2 differentiation pathway, instead the vast majority of cells remained GATA3^Lo^ Tbet^Lo^.

Furthermore, under these conditions, cells did not diverge into Treg [63] or Th17 [64] lineages. In cells activated with a strong TCR-signal the requirement for a third signal in order to differentiate to an effector cell likely acts as an additional checkpoint to inhibit aberrant Th1 responses and prevent spontaneous autoimmunity[65; 66]. Reinforcing this view, *in vivo* data have shown, that in response to a bacterial infection the absence of IL-12 reduces Th1 differentiation but does not lead to the default induction of Th2 differentiation [67]. Hence, in an *in vivo* polyclonal repertoire, CD4+ T cells responding to an activation stimulus in the absence of paracrine signal 3 inducing cytokines may exhibit significant Th2 skewing. Due to the potential for the activation of a large number of CD4+ T cells with comparatively weakly stimulated TCRs within a diverse repertoire. Under these conditions the absence of any repressive effect from signal 3 pathways would be permissive for Th2 differentiation. As seen in Milner *et al.* where the removal of high affinity cells from a polyclonal population led to the induction of Th2 differentiation in the remaining low affinity cells present [26]. Further, the priming of polyclonal naive cells with relatively low concentrations of peptide using a peptide library-based approach was also shown to preferentially induce Th2 skewing[68]. By comparison, in the presence of signal 3 inducing cytokines, CD4+ T cells encountering both weak and strong TCR signals would directly differentiate towards an alternate T-helper fate. As seen in the present study, where addition of exogenous IL-12 lead to a substantial polarization of the response towards Th1 differentiation, in a manner that synergized with strong TCR signalling and was largely dominant over weak TCR signalling.

Equally, the use of IL-4 at supraphysiological levels which has been classically used to drive *in vitro* Th2 differentiation[47] also dominated over TCR signal strength induced differentiation. Though it is important to note that in both cases of activation in the presence of endogenous cytokine stimuli, significant effects on GATA3 and Tbet expression levels driven by the strength of TCR signalling remained post-differentiation. Although, we observed both autocrine and exogenous IL-4 enhancement of Th2 differentiation, those cDC2 that have been shown to initiate *in vivo* Th2 differentiation [16; 17] do not provide an initial source of IL-4 to act as a polarizing signal 3 [13]. As such, it is possible that Th2 inducing cDC2 provide an initial TCR associated stimulatory signal that results in the preferential upregulation of GATA3, which is in turn enhanced by the production of autocrine IL-4. Supporting this theory are data from a recent study comparing LPS and papain driven *in vivo* T-helper responses, where it was shown that the cDC2 which drove the Th2 response presented considerably lower levels of cognate peptide in comparison to the Th1 inducing cDC2 [69]. Interestingly, these Th2 inducing cDC2 were also shown to exhibit a significant upregulation of costimulatory receptors, indicating the importance of signal 2 inputs for Th2 differentiation.

In the current study we have limited our scope to studying the effects of varying signal 1 during activation. However, in order to construct a unified model of differentiation the incorporation of the distinct effects of signal 2 inducing co-stimulatory molecules is also essential. As seen in previous studies, assessing the role of costimulation in modifying differentiation under *in vitro* conditions[69; 70; 71]. Recently, MARCH1 a ubiquitin ligase that regulates MHCII and CD86 turnover was identified as having a critical role for *in vivo* Th2 differentiation. As MARCH1 deficiency resulted in an increased accumulation of MHCII and CD86 on DCs and persistent T cell signalling, which induced increased Tbet and decreased GATA3 expression[72]. Our observations here, are also consistent with data from *in vivo* studies[26; 28; 73; 74] and those which have shown that nematode- and allergen-associated factors, such as omega-1 [75], Derp1 [76], and cystatins [77], downregulate signalling molecules in a way that alters the overall amount of TCR-associated signal received from activating APCs, either by altering p:MHCII levels, costimulatory expression (CD80/CD86), or DC morphology, to specifically drive a Th2 response. Reduced levels of TCR associated signalling may also be induced when activation occurs in the presence of DC surface markers previously associated with Th2 differentiation, such as OX40L [78], Notch [79], and CTLA-4. The biological relevance of this signalling strategy can be observed in responses to nematode parasites have developed a vast array of methodologies to evade the immune system. Through mechanisms including down regulation of CD3 expression[80], disruption of PAMP signalling[81; 82] and inhibition of IL-12[83] production. As such the Th2 response may have co-evolved without the requirement for a third activation signal in order to effectively counter these strategies[84].

At the population level, we initially observed that the strength of TCR-signalling altered mean expression levels of Tbet and GATA3. Based on subsequent mathematical modelling we determined that the ability of an individual cell to produce effector cytokines upon restimulation could be predicted as a stochastic function based on the ratiometric expression of GATA3 to Tbet. Studies describing the relationship between GATA3 and Tbet have previously shown these factors to be mutually antagonistic acting through several distinct mechanisms [4]. Such as Tbet binding and negatively regulating Th2-associated genes, including GATA3 [14]. Conversely, GATA3 impedes Th1 differentiation and Tbet expression through inhibitory actions on Runx3, STAT4, and IL-12rβ2 [85], and also represses the *Tbx21* locus through epigenetic modification [86]. GATA3-Tbet [87] and GATA3-Runx3 [85] protein-protein interactions have also been observed, whereby the competitive binding and sequestration of antagonistic master TFs results in suppression [88]. Together these observations provide a mechanistic rationale for our observations that the GATA3:Tbet level is a probabilistic determinant of cytokine expression and may account for the phenomena also noted in earlier studies [23; 24; 27], in which intermediate TCR signals lead to the inability of cells to produce either Th1- or Th2-effector cytokines upon restimulation. With regards to these states, probabilistic models have been described that account for the influence of TCR signal quality [89] and input from multiple cytokine signals [56; 57] on cell fate and even amongst strongly polarized cells, cytokine expression is largely a stochastic event, with only a limited proportion of cells expressing cytokines at any given time [58; 90].

In addition to TCR signal strength directing the outcome of differentiation, *in vivo* studies have shown the duration of cell contact and TCR signalling also affects Th-naïve activation [91; 92]. Our previous *in vivo* results [28] indicated that Th2 differentiation was associated with shorter periods of T cell–APC interaction, whereas stabilized interactions led to increased Th1 differentiation. Here, under *in vitro* conditions, T cell–APC interactions were limited to 2h in length, similar to those observed during phase I *in vivo* interactions [51]. We found that these initial interactions were sufficient to induce the signal-strength-dependent differentiation of *bona fide* Th1 and Th2 effector cells. Whilst previous studies had indicated cell division was an important component in the differentiation pathway [93; 94]. Instead, our data suggest that TCR signalling triggers two distinct cellular pathways, whereby initial contacts with an APC induces a TCR signal strength dependent priming for cell fate by promoting the expression of lineage-specific master TFs. Whereas the duration of TCR signalling directs entry into the cell cycle and is responsible for determining the number of rounds of cellular division. Similar results have previously been reported in the case of CD4+ T cells with mutated CD3 ITAM sequences [95], where a reduction in phosphorylatable ITAMs was permissive for IL-2 production but was insufficient to induce division. Further, Di Toro *et al.* [96] demonstrated that Tfh differentiation exhibits a similar TCR signal related bifurcation of outcomes, where strong TCR signalling preferentially induced BCL6 before the initiation of division, allowing for downstream Tfh differentiation. Conversely, weak signalling preferentially induced IL-2 responsiveness, upregulation of Blimp-1, and progression to non-Tfh fates.

Taken together our data establish a model of differentiation which accounts for many of the previously observed *in vivo* features of Th2 induction. Here, low levels of TCR stimuli directly induced Th2 differentiation independent of the need for a signal 3 cytokine input. Demonstrating how a quantitative signalling mechanism is permissive for the initial IL-4 independent induction of GATA3 that occurs during a type 2 inflammatory response and providing an explanation for the Th2 paradox. In contrast Th1 differentiation was shown to be dependent on presence of signal 3, either through autocrine IFNγ production induced by strong TCR signalling, or through the presence of an exogenous cytokine such as IL-12.

Mechanistically a bifurcation in T-helper differentiation pathways based on strength of activation stimuli may represent an additional checkpoint in the activation/differentiation cascade. Acting to prevent the unintentional induction of the more highly pathological Th1 and Th17[97; 98] responses, in the absence of the appropriate PAMP signals required for the induction of specific signal 3 cytokines, whilst enabling the detection of nematode parasites actively seeking to evade immune detection.

## Acknowledgements

We thank Dr Ronald Germain for his support during the study and for the kind provision of the B10.A 5C.C7 RAG2^−/−^ mice. Dr William E. Paul and Dr Hidehiro Yamane, for initial discussions and input into setting up the model system and Prof. Thomas Höfer for assistance with mathematical modelling approaches. The authors appreciate the help of Dr Vijay Govindharajan and LARC personnel for their assistance with animal husbandry. NVP was supported by NZFRST grant 20080148, TK was supported by QRDI grant NPRP11S-0122- 180359.

## Notes

### Competing Interest Statement

The authors have declared no competing interest.

## References

[1] C.S. Hsieh, S.E. Macatonia, C.S. Tripp, S.F. Wolf, A. O’Garra, and K.M. Murphy, Development of TH1 CD4+ T cells through IL-12 produced by Listeria-induced macrophages. Science 260 (1993) 547–9.

[2] P. Bocek, Jr., G. Foucras, and W.E. Paul, Interferon gamma enhances both in vitro and in vivo priming of CD4+ T cells for IL-4 production. The Journal of experimental medicine 199 (2004) 1619–30.

[3] G. Le Gros, S.Z. Ben-Sasson, R. Seder, F.D. Finkelman, and W.E. Paul, Generation of interleukin 4 (IL-4)-producing cells in vivo and in vitro: IL-2 and IL-4 are required for in vitro generation of IL-4-producing cells. The Journal of experimental medicine 172 (1990) 921–9.

[4] J. Zhu, and W.E. Paul, Peripheral CD4+ T-cell differentiation regulated by networks of cytokines and transcription factors. Immunological reviews 238 (2010) 247–62.

[5] C.A. Janeway, Jr., and R. Medzhitov, Innate immune recognition. Annu Rev Immunol 20 (2002) 197–216.

[6]. F.D. Finkelman, S.C. Morris, T. Orekhova, M. Mori, D. Donaldson, S.L. Reiner, N.L. Reilly, L. Schopf, and J.F. Urban, Jr., Stat6 regulation of in vivo IL-4 responses. Journal of immunology 164 (2000) 2303–10.

[7] M.H. Kaplan, A.L. Wurster, S.T. Smiley, and M.J. Grusby, Stat6-dependent and - independent pathways for IL-4 production. Journal of immunology 163 (1999) 6536–40.

[8] N. van Panhuys, S.-C. Tang, M. Prout, M. Camberis, D. Scarlett, J. Roberts, J. Hu-Li, W.E. Paul, and G. Le Gros, In vivo studies fail to reveal a role for IL-4 or STAT6 signaling in Th2 lymphocyte differentiation. Proceedings of the National Academy of Sciences of the United States of America 105 (2008) 12423–12428.

[9] B. Min, M. Prout, J. Hu-Li, J. Zhu, D. Jankovic, E.S. Morgan, J.F. Urban, Jr., A.M. Dvorak, F.D. Finkelman, G. LeGros, and W.E. Paul, Basophils Produce IL-4 and Accumulate in Tissues after Infection with a Th2-inducing Parasite. J. Exp. Med. %R 10.1084/jem.20040590 200 (2004) 507-517.

[10] D. Voehringer, K. Shinkai, and R.M. Locksley, Type 2 immunity reflects orchestrated recruitment of cells committed to IL-4 production. Immunity 20 (2004) 267–77.

[11] H. Hammad, and B.N. Lambrecht, Barrier Epithelial Cells and the Control of Type 2 Immunity. Immunity 43 (2015) 29–40.

[12] S.A. Islam, and A.D. Luster, T cell homing to epithelial barriers in allergic disease. Nature medicine 18 (2012) 705–15.

[13] B. Leon, A model of Th2 differentiation based on polarizing cytokine repression. Trends Immunol 44 (2023) 399–407.

[14] J. Zhu, D. Jankovic, A.J. Oler, G. Wei, S. Sharma, G. Hu, L. Guo, R. Yagi, H. Yamane, G. Punkosdy, L. Feigenbaum, K. Zhao, and W.E. Paul, The transcription factor T-bet is induced by multiple pathways and prevents an endogenous Th2 cell program during Th1 cell responses. Immunity 37 (2012) 660–73.

[15] D. Jankovic, S. Steinfelder, M.C. Kullberg, and A. Sher, Mechanisms underlying helminth- induced Th2 polarization: default, negative or positive pathways? Chem Immunol Allergy 90 (2006) 65–81.

[16] M. Plantinga, M. Guilliams, M. Vanheerswynghels, K. Deswarte, F. Branco-Madeira, W. Toussaint, L. Vanhoutte, K. Neyt, N. Killeen, B. Malissen, H. Hammad, and B.N. Lambrecht, Conventional and monocyte-derived CD11b(+) dendritic cells initiate and maintain T helper 2 cell-mediated immunity to house dust mite allergen. Immunity 38 (2013) 322–35.

[17] J.U. Mayer, K.L. Hilligan, J.S. Chandler, D.A. Eccles, S.I. Old, R.G. Domingues, J. Yang, G.R. Webb, L. Munoz-Erazo, E.J. Hyde, K.A. Wakelin, S.C. Tang, S.C. Chappell, S. von Daake, F. Brombacher, C.R. Mackay, A. Sher, R. Tussiwand, L.M. Connor, D. Gallego-Ortega, D. Jankovic, G. Le Gros, M.R. Hepworth, O. Lamiable, and F. Ronchese, Homeostatic IL-13 in healthy skin directs dendritic cell differentiation to promote T(H)2 and inhibit T(H)17 cell polarization. Nature immunology 22 (2021) 1538–1550.

[18] Y. Gao, S.A. Nish, R. Jiang, L. Hou, P. Licona-Limon, J.S. Weinstein, H. Zhao, and R. Medzhitov, Control of T helper 2 responses by transcription factor IRF4-dependent dendritic cells. Immunity 39 (2013) 722–32.

[19] R. Tussiwand, B. Everts, G.E. Grajales-Reyes, N.M. Kretzer, A. Iwata, J. Bagaitkar, X. Wu, R. Wong, D.A. Anderson, T.L. Murphy, E.J. Pearce, and K.M. Murphy, Klf4 expression in conventional dendritic cells is required for T helper 2 cell responses. Immunity 42 (2015) 916–28.

[20] T. Yoshimoto, The Hunt for the Source of Primary Interleukin-4: How We Discovered That Natural Killer T Cells and Basophils Determine T Helper Type 2 Cell Differentiation In Vivo. Front Immunol 9 (2018) 716.

[21] S. Kumar, Y. Jeong, M.U. Ashraf, and Y.S. Bae, Dendritic Cell-Mediated Th2 Immunity and Immune Disorders. Int J Mol Sci 20 (2019).

[22] N. Noben-Trauth, J. Hu-Li, and W.E. Paul, IL-4 secreted from individual naive CD4+ T cells acts in an autocrine manner to induce Th2 differentiation. Eur J Immunol 32 (2002) 1428–33.

[23] J.L. Brogdon, D. Leitenberg, and K. Bottomly, The potency of TCR signaling differentially regulates NFATc/p activity and early IL-4 transcription in naive CD4+ T cells. Journal of immunology 168 (2002) 3825–32.

[24] S. Constant, C. Pfeiffer, A. Woodard, T. Pasqualini, and K. Bottomly, Extent of T cell receptor ligation can determine the functional differentiation of naive CD4+ T cells. The Journal of experimental medicine 182 (1995) 1591–6.

[25] P.J. Jorritsma, J.L. Brogdon, and K. Bottomly, Role of TCR-induced extracellular signal-regulated kinase activation in the regulation of early IL-4 expression in naive CD4+ T cells. Journal of immunology 170 (2003) 2427–34.

[26] J.D. Milner, N. Fazilleau, M. McHeyzer-Williams, and W. Paul, Cutting edge: lack of high affinity competition for peptide in polyclonal CD4+ responses unmasks IL-4 production. Journal of immunology 184 (2010) 6569–73.

[27] H. Yamane, J. Zhu, and W.E. Paul, Independent roles for IL-2 and GATA-3 in stimulating naive CD4+ T cells to generate a Th2-inducing cytokine environment. J. Exp. Med. 202 (2005) 793–804.

[28] N. van Panhuys, F. Klauschen, and R.N. Germain, T-Cell-Receptor-Dependent Signal Intensity Dominantly Controls CD4(+) T Cell Polarization In Vivo. Immunity 41 (2014) 63–74.

[29] J. Pulecio, J. Petrovic, F. Prete, G. Chiaruttini, A.M. Lennon-Dumenil, C. Desdouets, S. Gasman, O.R. Burrone, and F. Benvenuti, Cdc42-mediated MTOC polarization in dendritic cells controls targeted delivery of cytokines at the immune synapse. The Journal of experimental medicine 207 (2010) 2719–32.

[30] R.A. Maldonado, D.J. Irvine, R. Schreiber, and L.H. Glimcher, A role for the immunological synapse in lineage commitment of CD4 lymphocytes. Nature 431 (2004) 527–32.

[31] M. Huse, B.F. Lillemeier, M.S. Kuhns, D.S. Chen, and M.M. Davis, T cells use two directionally distinct pathways for cytokine secretion. Nature immunology 7 (2006) 247–55.

[32] L. Ding, P.S. Linsley, L.Y. Huang, R.N. Germain, and E.M. Shevach, IL-10 inhibits macrophage costimulatory activity by selectively inhibiting the up-regulation of B7 expression. Journal of immunology 151 (1993) 1224–34.

[33] M.K.R. Kalikiri, H.S. Manjunath, F.R. Vempalli, L.S. Mathew, L. Liu, L. Wang, G. Wang, K. Wang, O. Soloviov, S. Lorenz, and S. Tomei, Technical assessment of different extraction methods and transcriptome profiling of RNA isolated from small volumes of blood. Sci Rep 13 (2023) 3598.

[34] S. Anders, and W. Huber, Differential expression analysis for sequence count data. Genome Biol 11 (2010) R106.

[35] S.X. Ge, E.W. Son, and R. Yao, iDEP: an integrated web application for differential expression and pathway analysis of RNA-Seq data. BMC Bioinformatics 19 (2018) 534.

[36] A. Subramanian, P. Tamayo, V.K. Mootha, S. Mukherjee, B.L. Ebert, M.A. Gillette, A. Paulovich, S.L. Pomeroy, T.R. Golub, E.S. Lander, and J.P. Mesirov, Gene set enrichment analysis: a knowledge-based approach for interpreting genome-wide expression profiles. Proc Natl Acad Sci U S A 102 (2005) 15545–50.

[37] D. Chauss, T. Freiwald, R. McGregor, B. Yan, L. Wang, E. Nova-Lamperti, D. Kumar, Z. Zhang, H. Teague, E.E. West, K.M. Vannella, M.J. Ramos-Benitez, J. Bibby, A. Kelly, A. Malik, A.F. Freeman, D.M. Schwartz, D. Portilla, D.S. Chertow, S. John, P. Lavender, C. Kemper, G. Lombardi, N.N. Mehta, N. Cooper, M.S. Lionakis, A. Laurence, M. Kazemian, and B. Afzali, Autocrine vitamin D signaling switches off pro-inflammatory programs of T(H)1 cells. Nature immunology 23 (2022) 62–74.

[38] M.J. Stubbington, B. Mahata, V. Svensson, A. Deonarine, J.K. Nissen, A.G. Betz, and S.A. Teichmann, An atlas of mouse CD4(+) T cell transcriptomes. Biol Direct 10 (2015) 14.

[39] L.J. Gearing, H.E. Cumming, R. Chapman, A.M. Finkel, I.B. Woodhouse, K. Luu, J.A. Gould, S.C. Forster, and P.J. Hertzog, CiiiDER: A tool for predicting and analysing transcription factor binding sites. PloS one 14 (2019) e0215495.

[40] J. Xia, N. Psychogios, N. Young, and D.S. Wishart, MetaboAnalyst: a web server for metabolomic data analysis and interpretation. Nucleic Acids Res 37 (2009) W652–60.

[41] J.M. Elizarraras, Y. Liao, Z. Shi, Q. Zhu, A.R. Pico, and B. Zhang, WebGestalt 2024: faster gene set analysis and new support for metabolomics and multi-omics. Nucleic Acids Res 52 (2024) W415–W421.

[42] C. Helmstetter, M. Flossdorf, M. Peine, A. Kupz, J. Zhu, A.N. Hegazy, M.A. Duque-Correa, Q. Zhang, Y. Vainshtein, A. Radbruch, S.H. Kaufmann, W.E. Paul, T. Hofer, and M. Lohning, Individual T helper cells have a quantitative cytokine memory. Immunity 42 (2015) 108–22.

[43] J. Hu-Li, C. Pannetier, L. Guo, M. Lohning, H. Gu, C. Watson, M. Assenmacher, A. Radbruch, and W.E. Paul, Regulation of expression of IL-4 alleles: analysis using a chimeric GFP/IL-4 gene. Immunity 14 (2001) 1–11.

[44] K.A. Allison, E. Sajti, J.G. Collier, D. Gosselin, T.D. Troutman, E.L. Stone, S.M. Hedrick, and C.K. Glass, Affinity and dose of TCR engagement yield proportional enhancer and gene activity in CD4+ T cells. Elife 5 (2016).

[45] S.J. Szabo, S.T. Kim, G.L. Costa, X. Zhang, C.G. Fathman, and L.H. Glimcher, A novel transcription factor, T-bet, directs Th1 lineage commitment. Cell 100 (2000) 655–69.

[46] H. Ariga, Y. Shimohakamada, M. Nakada, T. Tokunaga, T. Kikuchi, A. Kariyone, T. Tamura, and K. Takatsu, Instruction of naive CD4+ T-cell fate to T-bet expression and T helper 1 development: roles of T-cell receptor-mediated signals. Immunology 122 (2007) 210–21.

[47] M.A. Al-Aghbar, M. Espino Guarch, and N. van Panhuys, IL-2 amplifies quantitative TCR signalling inputs to drive Th1 and Th2 differentiation. Immunology (2024).

[48] W. Zheng, and R.A. Flavell, The transcription factor GATA-3 is necessary and sufficient for Th2 cytokine gene expression in CD4 T cells. Cell 89 (1997) 587–96.

[49] J. Zhu, H. Yamane, J. Cote-Sierra, L. Guo, and W.E. Paul, GATA-3 promotes Th2 responses through three different mechanisms: induction of Th2 cytokine production, selective growth of Th2 cells and inhibition of Th1 cell-specific factors. Cell Res 16 (2006) 3–10.

[50] P. Burt, M. Peine, C. Peine, Z. Borek, S. Serve, M. Flossdorf, A.N. Hegazy, T. Hofer, M. Lohning, and K. Thurley, Dissecting the dynamic transcriptional landscape of early T helper cell differentiation into Th1, Th2, and Th1/2 hybrid cells. Front Immunol 13 (2022) 928018.

[51] S.E. Henrickson, T.R. Mempel, I.B. Mazo, B. Liu, M.N. Artyomov, H. Zheng, A. Peixoto, M.P. Flynn, B. Senman, T. Junt, H.C. Wong, A.K. Chakraborty, and U.H. von Andrian, T cell sensing of antigen dose governs interactive behavior with dendritic cells and sets a threshold for T cell activation. Nature immunology 9 (2008) 282–91.

[52] J.B. Huppa, M. Gleimer, C. Sumen, and M.M. Davis, Continuous T cell receptor signaling required for synapse maintenance and full effector potential. Nature immunology 4 (2003) 749–55.

[53] J. Zhu, H. Yamane, and W.E. Paul, Differentiation of effector CD4 T cell populations (*). Annu Rev Immunol 28 (2010) 445–89.

[54] J. Zhu, T Helper Cell Differentiation, Heterogeneity, and Plasticity. Cold Spring Harb Perspect Biol 10 (2018).

[55] L. Zhou, M.M. Chong, and D.R. Littman, Plasticity of CD4+ T cell lineage differentiation. Immunity 30 (2009) 646–55.

[56] B.L. Puniya, R.G. Todd, A. Mohammed, D.M. Brown, M. Barberis, and T. Helikar, A Mechanistic Computational Model Reveals That Plasticity of CD4(+) T Cell Differentiation Is a Function of Cytokine Composition and Dosage. Front Physiol 9 (2018) 878.

[57] I. Eizenberg-Magar, J. Rimer, I. Zaretsky, D. Lara-Astiaso, S. Reich-Zeliger, and N. Friedman, Diverse continuum of CD4(+) T-cell states is determined by hierarchical additive integration of cytokine signals. Proc Natl Acad Sci U S A 114 (2017) E6447–E6456.

[58] M. Peine, S. Rausch, C. Helmstetter, A. Frohlich, A.N. Hegazy, A.A. Kuhl, C.G. Grevelding, T. Hofer, S. Hartmann, and M. Lohning, Stable T-bet(+)GATA-3(+) Th1/Th2 hybrid cells arise in vivo, can develop directly from naive precursors, and limit immunopathologic inflammation. PLoS Biol 11 (2013) e1001633.

[59]. A. Das, V. Ranganathan, D. Umar, S. Thukral, A. George, S. Rath, and V. Bal, Effector/memory CD4 T cells making either Th1 or Th2 cytokines commonly co-express T-bet and GATA-3. PloS one 12 (2017) e0185932.

[60] Y.E. Antebi, S. Reich-Zeliger, Y. Hart, A. Mayo, I. Eizenberg, J. Rimer, P. Putheti, D. Pe’er, and N. Friedman, Mapping differentiation under mixed culture conditions reveals a tunable continuum of T cell fates. PLoS Biol 11 (2013) e1001616.

[61] F. Yu, S. Sharma, J. Edwards, L. Feigenbaum, and J. Zhu, Dynamic expression of transcription factors T-bet and GATA-3 by regulatory T cells maintains immunotolerance. Nature immunology 16 (2015) 197–206.

[62] E.A. Wohlfert, J.R. Grainger, N. Bouladoux, J.E. Konkel, G. Oldenhove, C.H. Ribeiro, J.A. Hall, R. Yagi, S. Naik, R. Bhairavabhotla, W.E. Paul, R. Bosselut, G. Wei, K. Zhao, M. Oukka, J. Zhu, and Y. Belkaid, GATA3 controls Foxp3(+) regulatory T cell fate during inflammation in mice. J Clin Invest 121 (2011) 4503–15.

[63] R.A. Gottschalk, E. Corse, and J.P. Allison, TCR ligand density and affinity determine peripheral induction of Foxp3 in vivo. The Journal of experimental medicine 207 (2010) 1701–11.

[64] I. Kastirr, M. Crosti, S. Maglie, M. Paroni, B. Steckel, M. Moro, M. Pagani, S. Abrignani, and J. Geginat, Signal Strength and Metabolic Requirements Control Cytokine-Induced Th17 Differentiation of Uncommitted Human T Cells. Journal of immunology 195 (2015) 3617–27.

[65] J.F. Ashouri, W.L. Lo, T.T.T. Nguyen, L. Shen, and A. Weiss, ZAP70, too little, too much can lead to autoimmunity. Immunological reviews 307 (2022) 145–160.

[66] H. Jiang, and L. Chess, How the immune system achieves self-nonself discrimination during adaptive immunity. Advances in immunology 102 (2009) 95–133.

[67] D. Jankovic, M.C. Kullberg, S. Hieny, P. Caspar, C.M. Collazo, and A. Sher, In the absence of IL-12, CD4(+) T cell responses to intracellular pathogens fail to default to a Th2 pattern and are host protective in an IL-10(-/-) setting. Immunity 16 (2002) 429–39.

[68] J.S. Barber, L.K. Yokomizo, V. Sheikh, A.F. Freeman, E. Garabedian, E. van Dijk, R. Sokolic, F. Candotti, N.P. Weng, I. Sereti, and J.D. Milner, Peptide library-based evaluation of T-cell receptor breadth detects defects in global and regulatory activation in human immunologic diseases. Proc Natl Acad Sci U S A 110 (2013) 8164–9.

[69] M.R. Lyons-Cohen, E.A. Shamskhou, and M.Y. Gerner, Site-specific regulation of Th2 differentiation within lymph node microenvironments. The Journal of experimental medicine 221 (2024).

[70] P.R. Rogers, and M. Croft, CD28, Ox-40, LFA-1, and CD4 modulation of Th1/Th2 differentiation is directly dependent on the dose of antigen. Journal of immunology 164 (2000) 2955–63.

[71] X. Tao, S. Constant, P. Jorritsma, and K. Bottomly, Strength of TCR signal determines the costimulatory requirements for Th1 and Th2 CD4+ T cell differentiation. Journal of immunology 159 (1997) 5956–63.

[72] C.A. Castellanos, X. Ren, S.L. Gonzalez, H.K. Li, A.W. Schroeder, H.E. Liang, B.J. Laidlaw, D. Hu, A.C.Y. Mak, C. Eng, J.R. Rodriguez-Santana, M. LeNoir, Q. Yan, J.C. Celedon, E.G. Burchard, S.S. Zamvil, S. Ishido, R.M. Locksley, J.G. Cyster, X. Huang, and J.S. Shin, Lymph node-resident dendritic cells drive T(H)2 cell development involving MARCH1. Sci Immunol 6 (2021) eabh0707.

[73] B.J. Marsland, T.J. Soos, G. Spath, D.R. Littman, and M. Kopf, Protein kinase C theta is critical for the development of in vivo T helper (Th)2 cell but not Th1 cell responses. The Journal of experimental medicine 200 (2004) 181–9.

[74] O.M. Siggs, L.A. Miosge, A.L. Yates, E.M. Kucharska, D. Sheahan, T. Brdicka, A. Weiss, A. Liston, and C.C. Goodnow, Opposing functions of the T cell receptor kinase ZAP-70 in immunity and tolerance differentially titrate in response to nucleotide substitutions. Immunity 27 (2007) 912–26.

[75] S. Steinfelder, J.F. Andersen, J.L. Cannons, C.G. Feng, M. Joshi, D. Dwyer, P. Caspar, P.L. Schwartzberg, A. Sher, and D. Jankovic, The major component in schistosome eggs responsible for conditioning dendritic cells for Th2 polarization is a T2 ribonuclease (omega-1). The Journal of experimental medicine 206 (2009) 1681–90.

[76] R. Furmonaviciene, A.M. Ghaemmaghami, S.E. Boyd, N.S. Jones, K. Bailey, A.C. Willis, H.F. Sewell, D.A. Mitchell, and F. Shakib, The protease allergen Der p 1 cleaves cell surface DC-SIGN and DC-SIGNR: experimental analysis of in silico substrate identification and implications in allergic responses. Clin Exp Allergy 37 (2007) 231–42.

[77] S. Hartmann, and R. Lucius, Modulation of host immune responses by nematode cystatins. Int J Parasitol 33 (2003) 1291–302.

[78] T. Ito, Y.H. Wang, O. Duramad, T. Hori, G.J. Delespesse, N. Watanabe, F.X. Qin, Z. Yao, W. Cao, and Y.J. Liu, TSLP-activated dendritic cells induce an inflammatory T helper type 2 cell response through OX40 ligand. The Journal of experimental medicine 202 (2005) 1213–23.

[79] H. Yamane, and W.E. Paul, Early signaling events that underlie fate decisions of naive CD4(+) T cells toward distinct T-helper cell subsets. Immunological reviews 252 (2013) 12–23.

[80] L.J. Appleby, N. Nausch, F. Heard, L. Erskine, C.D. Bourke, N. Midzi, T. Mduluza, J.E. Allen, and F. Mutapi, Down Regulation of the TCR Complex CD3zeta-Chain on CD3+ T Cells: A Potential Mechanism for Helminth-Mediated Immune Modulation. Front Immunol 6 (2015) 51.

[81] M. Segura, Z. Su, C. Piccirillo, and M.M. Stevenson, Impairment of dendritic cell function by excretory-secretory products: a potential mechanism for nematode-induced immunosuppression. Eur J Immunol 37 (2007) 1887–904.

[82] S. Rajasekaran, R. Anuradha, and R. Bethunaickan, TLR Specific Immune Responses against Helminth Infections. J Parasitol Res 2017 (2017) 6865789.

[83] A. Balic, Y. Harcus, M.J. Holland, and R.M. Maizels, Selective maturation of dendritic cells by Nippostrongylus brasiliensis-secreted proteins drives Th2 immune responses. Eur J Immunol 34 (2004) 3047–59.

[84] J.E. Allen, and T.E. Sutherland, Host protective roles of type 2 immunity: parasite killing and tissue repair, flip sides of the same coin. Semin Immunol 26 (2014) 329–40.

[85] R. Yagi, I.S. Junttila, G. Wei, J.F. Urban, Jr., K. Zhao, W.E. Paul, and J. Zhu, The transcription factor GATA3 actively represses RUNX3 protein-regulated production of interferon-gamma. Immunity 32 (2010) 507–17.

[86] G. Wei, B.J. Abraham, R. Yagi, R. Jothi, K. Cui, S. Sharma, L. Narlikar, D.L. Northrup, Q. Tang, W.E. Paul, J. Zhu, and K. Zhao, Genome-wide analyses of transcription factor GATA3-mediated gene regulation in distinct T cell types. Immunity 35 (2011) 299–311.

[87] E.S. Hwang, S.J. Szabo, P.L. Schwartzberg, and L.H. Glimcher, T Helper Cell Fate Specified by Kinase-Mediated Interaction of T-bet with GATA-3. Science 307 (2005) 430–433.

[88] A. Hertweck, M. Vila de Mucha, P.R. Barber, R. Dagil, H. Porter, A. Ramos, G.M. Lord, and R.G. Jenner, The TH1 cell lineage-determining transcription factor T-bet suppresses TH2 gene expression by redistributing GATA3 away from TH2 genes. Nucleic Acids Res 50 (2022) 4557–4573.

[89] Y.L. Cho, M. Flossdorf, L. Kretschmer, T. Hofer, D.H. Busch, and V.R. Buchholz, TCR Signal Quality Modulates Fate Decisions of Single CD4(+) T Cells in a Probabilistic Manner. Cell Rep 20 (2017) 806–818.

[90] L. Guo, J. Hu-Li, and W.E. Paul, Probabilistic Regulation in TH2 Cells Accounts for Monoallelic Expression of IL-4 and IL-13. Immunity 23 (2005) 89–99.

[91] S.E. Henrickson, T.R. Mempel, I.B. Mazo, B. Liu, M.N. Artyomov, H. Zheng, A. Peixoto, M. Flynn, B. Senman, T. Junt, H.C. Wong, A.K. Chakraborty, and U.H. von Andrian, In vivo imaging of T cell priming. Sci Signal 1 (2008) pt2.

[92] M.D. Cahalan, and I. Parker, Choreography of cell motility and interaction dynamics imaged by two-photon microscopy in lymphoid organs. Annu Rev Immunol 26 (2008) 585–626.

[93] A.V. Gett, and P.D. Hodgkin, Cell division regulates the T cell cytokine repertoire, revealing a mechanism underlying immune class regulation. Proc Natl Acad Sci U S A 95 (1998) 9488–93.

[94] R. Obst, The Timing of T Cell Priming and Cycling. Front Immunol 6 (2015) 563.

[95] C.S. Guy, K.M. Vignali, J. Temirov, M.L. Bettini, A.E. Overacre, M. Smeltzer, H. Zhang, J.B. Huppa, Y.H. Tsai, C. Lobry, J. Xie, P.J. Dempsey, H.C. Crawford, I. Aifantis, M.M. Davis, and D.A. Vignali, Distinct TCR signaling pathways drive proliferation and cytokine production in T cells. Nature immunology 14 (2013) 262–70.

[96] D. DiToro, C.J. Winstead, D. Pham, S. Witte, R. Andargachew, J.R. Singer, C.G. Wilson, C.L. Zindl, R.J. Luther, D.J. Silberger, B.T. Weaver, E.M. Kolawole, R.J. Martinez, H. Turner, R.D. Hatton, J.J. Moon, S.S. Way, B.D. Evavold, and C.T. Weaver, Differential IL-2 expression defines developmental fates of follicular versus nonfollicular helper T cells. Science 361 (2018).

[97] M. Mbow, B.M. Larkin, L. Meurs, L.J. Wammes, S.E. de Jong, L.A. Labuda, M. Camara, H.H. Smits, K. Polman, T.N. Dieye, S. Mboup, M.J. Stadecker, and M. Yazdanbakhsh, T-helper 17 cells are associated with pathology in human schistosomiasis. J Infect Dis 207 (2013) 186–95.

[98] S. Babu, S.Q. Bhat, N. Pavan Kumar, A.B. Lipira, S. Kumar, C. Karthik, V. Kumaraswami, and T.B. Nutman, Filarial lymphedema is characterized by antigen-specific Th1 and th17 proinflammatory responses and a lack of regulatory T cells. PLoS Negl Trop Dis 3 (2009) e420.

