## Supplemental Figures S1-S9 for "Initial TCR Signal Strength Imprints GATA3 and Tbet Expression Driving T-helper Cell Fate Decisions"

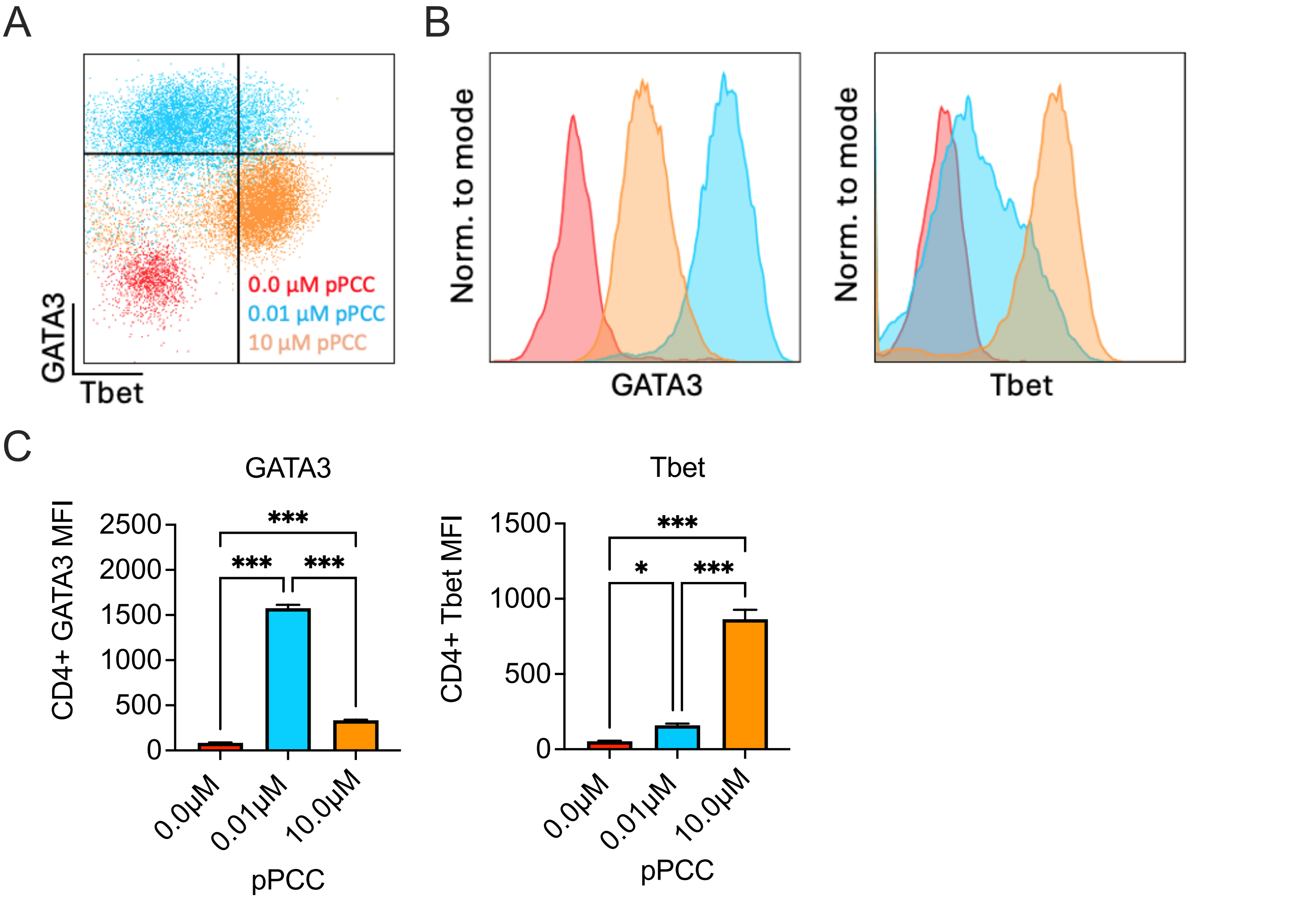


**Figure S1: Upregulation of GATA3 and Tbet in naïve compared to differentiated CD4+ T cells**

Naive 5CC7 CD4$^+$ T cells were stimulated with P13.9 antigen-presenting cells incubated with 0.0 µM (naive), 0.01 µM, or 10 µM pigeon cytochrome C peptide (pPCC) for 4 days under in vitro conditions. (A) Tbet vs. GATA3 expression was determined following intracellular staining. (B) Histogram comparison of normalized GATA3 and Tbet expression. (C) CD4+ T cell MFIs of GATA3 and Tbet expression. Error bars indicate mean ±SEM, n = 3; experiments were performed at least three times with consistent results. Statistical analysis was performed via 1-way ANOVA with Tukey multiple comparison testing. * = $p < 0.05$, ** = $p < 0.01$, *** = $p < 0.001$, ns = non-significant.

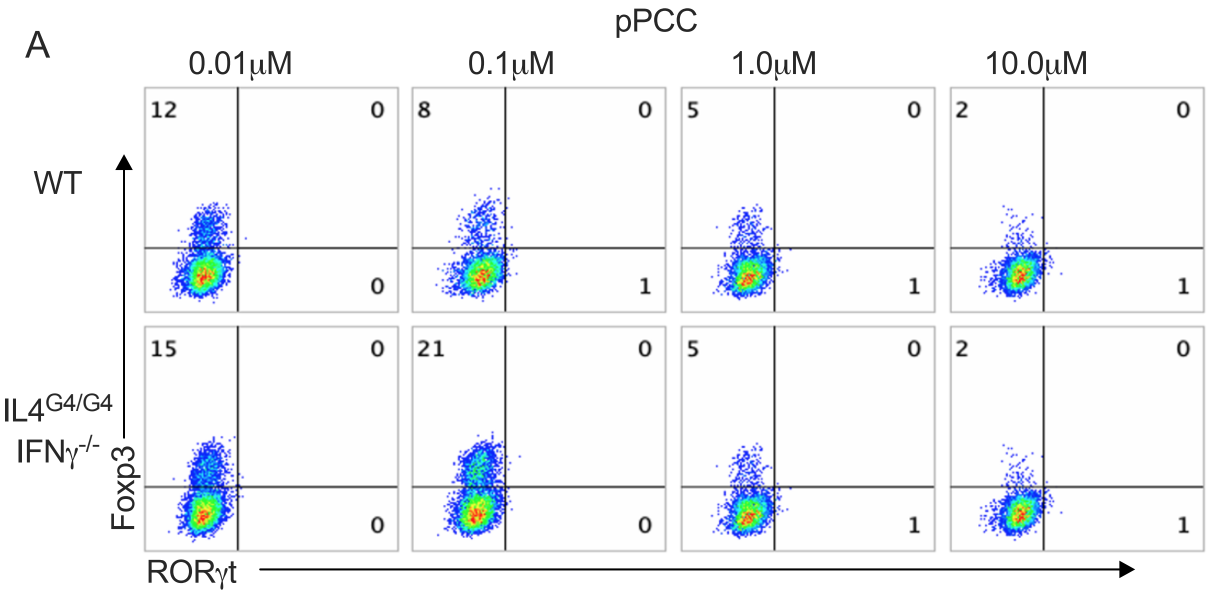


**Figure S2: Initial TCR stimulation induces Foxp3 expression but is not sufficient to induce RORγt expression.** (A) Naïve WT and IL-4$^{G4/G4}$ IFNγ$^{-/-}$ CD4$^{+}$ T cells were stimulated with APC, in the presence of 0.01–10 µM pPCC, for four days. Cells were then restimulated and the frequency of Foxp3 and RORγt was calculated following intracellular staining. Experiments were performed twice with consistent results.

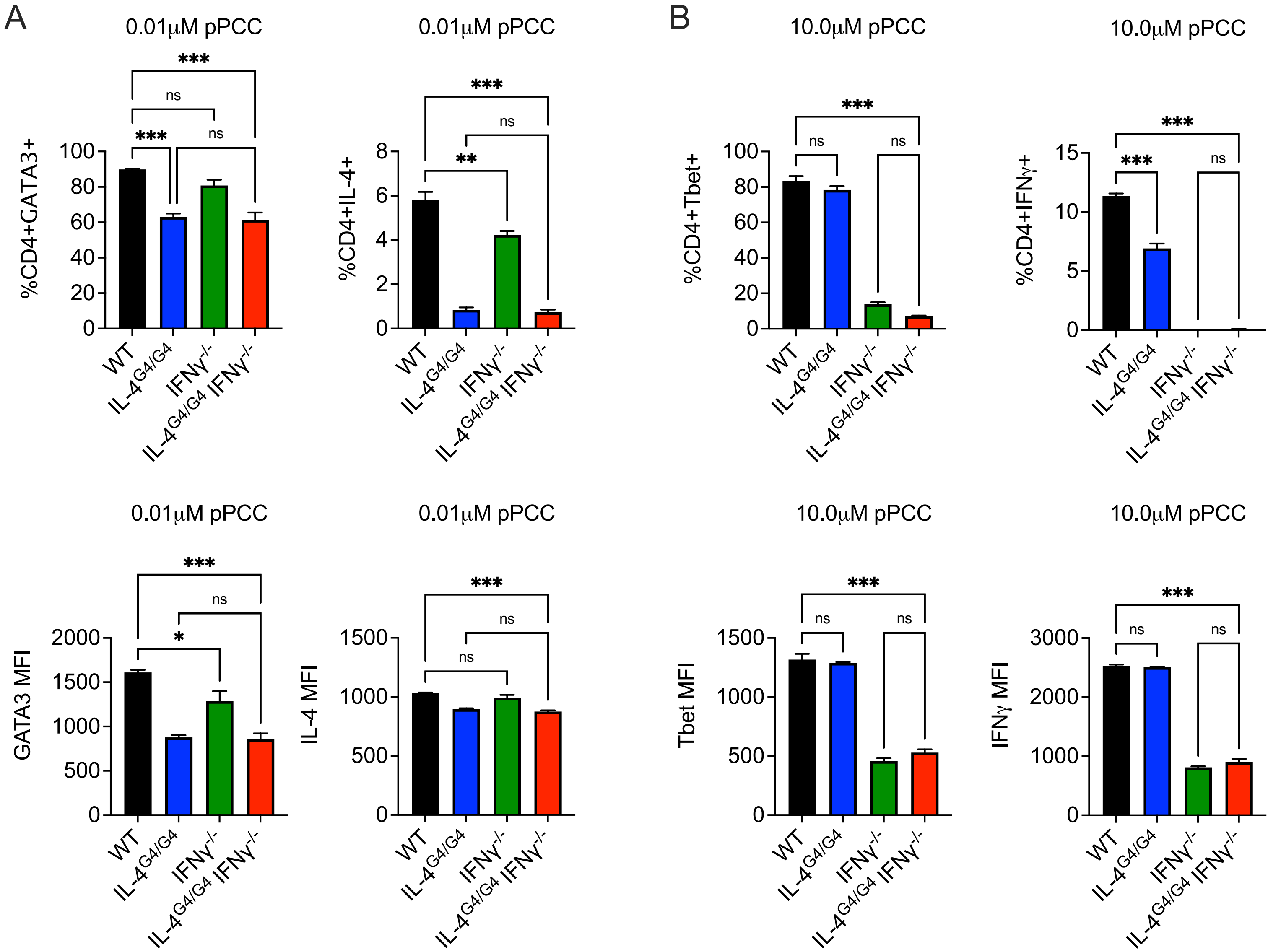


**Figure S3: IL-4 is not required for the induction of GATA3, IFNγ induces Tbet expression**

Naïve 5CC7 $CD4^{+}$ T cells from WT, IL-$4^{G4/G4}$, IFNγ$^{-/-}$, and IL-$4^{G4/G4}$ IFNγ$^{-/-}$ mice were stimulated with P13.9 antigen presenting cells in the presence of 0.01-10 μM pigeon cytochrome C peptide (pPCC) for 4 days under in vitro conditions. Cells were restimulated with PMA and ionomycin. IL-4, IFNγ, GATA3, and Tbet expression was determined by intracellular staining. (A) For cells stimulated with 0.01 μM pPCC, the frequencies and MFIs of GATA3 and IL-4 expression are shown. (B) For cells stimulated with 10.0 μM pPCC, the frequencies and MFIs of Tbet and IFNγ expression are shown. Error bars in (A&B) indicate mean ±SEM, n=3; experiments were performed at least three times with consistent results. Statistical analysis was performed via 1-way ANOVA with Tukey multiple comparison testing. * = $p < 0.05$, ** = $p < 0.01$, *** = $p < 0.001$, ns = non-significant.

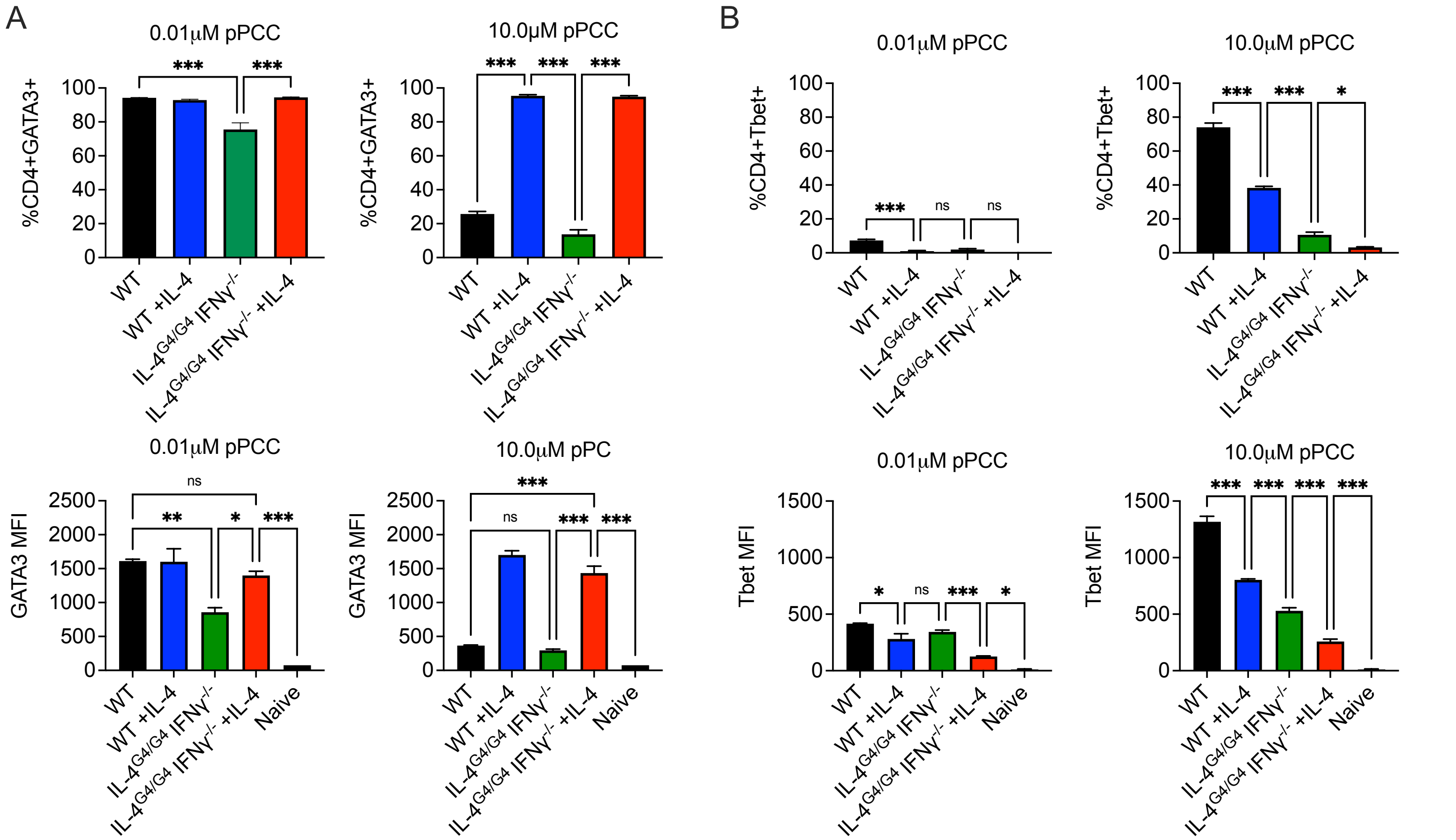


**Figure S4: IL-4 enhances GATA3 expression, and partially represses Tbet expression under Th1 inducing conditions**

Naïve 5CC7 CD4$^{+}$ T cells from WT and IL-4$^{G4/G4}$IFNγ$^{-/-}$ mice were stimulated in the presence of 0.01-10 μM pPCC, and exogenous IL-4 was added as indicated. (A) For cells stimulated with 0.01 μM and 10.0 μM pPCC, the frequencies and MFIs of GATA3 (A) and Tbet (B) expression are shown. Error bars indicate mean ±SEM, n = 3; experiments were performed at least three times with consistent results. Statistical analysis was performed via 1-way ANOVA with Tukey multiple comparison testing. * = $p < 0.05$, ** = $p < 0.01$, *** = $p < 0.001$, ns = non-significant.

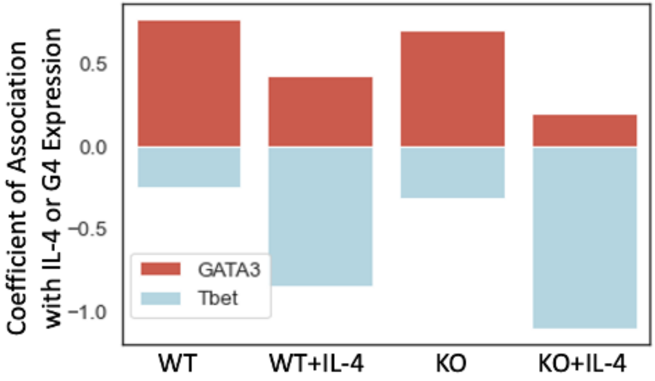


| Features | GATA3 | | Tbet | |
|---|---|---|---|---|
| Test | Coef. | nLog p Value (adjusted) | Coef. | nLog p Value (adjusted) |
| WT | 0.77 | 5.43 | -0.24 | <0.001 |
| WT + IL-4 | 0.42 | 1.58 | -0.84 | 7.73 |
| KO | 0.7 | 1.73 | -0.31 | <0.001 |
| KO + IL-4 | 0.19 | <0.001 | -1.11 | 17.97 |

**Figure S5: Relative contributions of transcription factors GATA3 and Tbet to coefficient of association for IL-4 expression**

The relative expression of IL-4 by wildtype (WT) 5CC7 cells or G4 expression by IL-4$^{G4/G4}$IFNγ$^{-/-}$ knockout (KO) cells in the presence of normalized GATA3 and Tbet expression in each condition was tested with a robust linear model, and the coefficients (Coef.) of association were plotted simultaneously for both transcription factors.

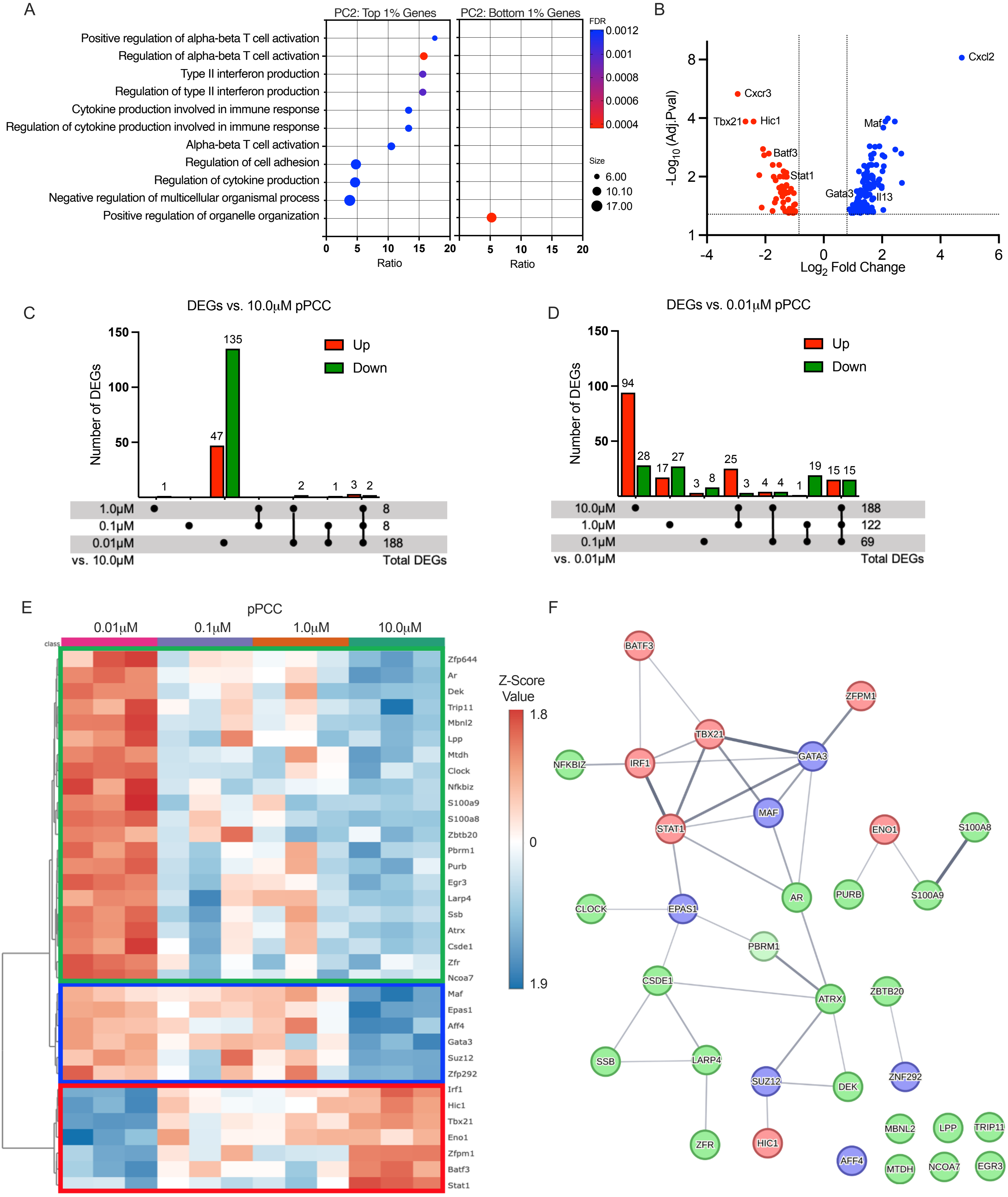


**Figure S6: Transcriptional analysis reveals graded expression of key regulatory modules induced during differentiation** (A) Gene ontology analysis of top and bottom 1% of genes identified as components of PC2 in PLSDA of Th-naïve and activated CD4+ T cells. (B) Volcano plot of DEGs from comparison of CD4+ T cells stimulated 10.0μM (Red) versus 0.01μM (Blue) pPCC. Numbers of differentially expressed genes in comparison of 10.0μM (C) and 0.01 μM (D) pPCC-activated CD4+ T cells to conditions indicated. (E) Transcription factors (TFs) present in all DEGs were identified, and expression levels present in individual samples used to generate a heatmap. Unbiased clustering of DEGs identified two major subsets of T-helper (Th)-associated genes Th1 (Red) and Th2 (Blue), with remaining genes shown in green. (F) TFs identified were used generate a STRING interaction network diagram with nodes coloured as in (E); intensity of edges indicates interaction confidence based on known functional and physical protein associations.

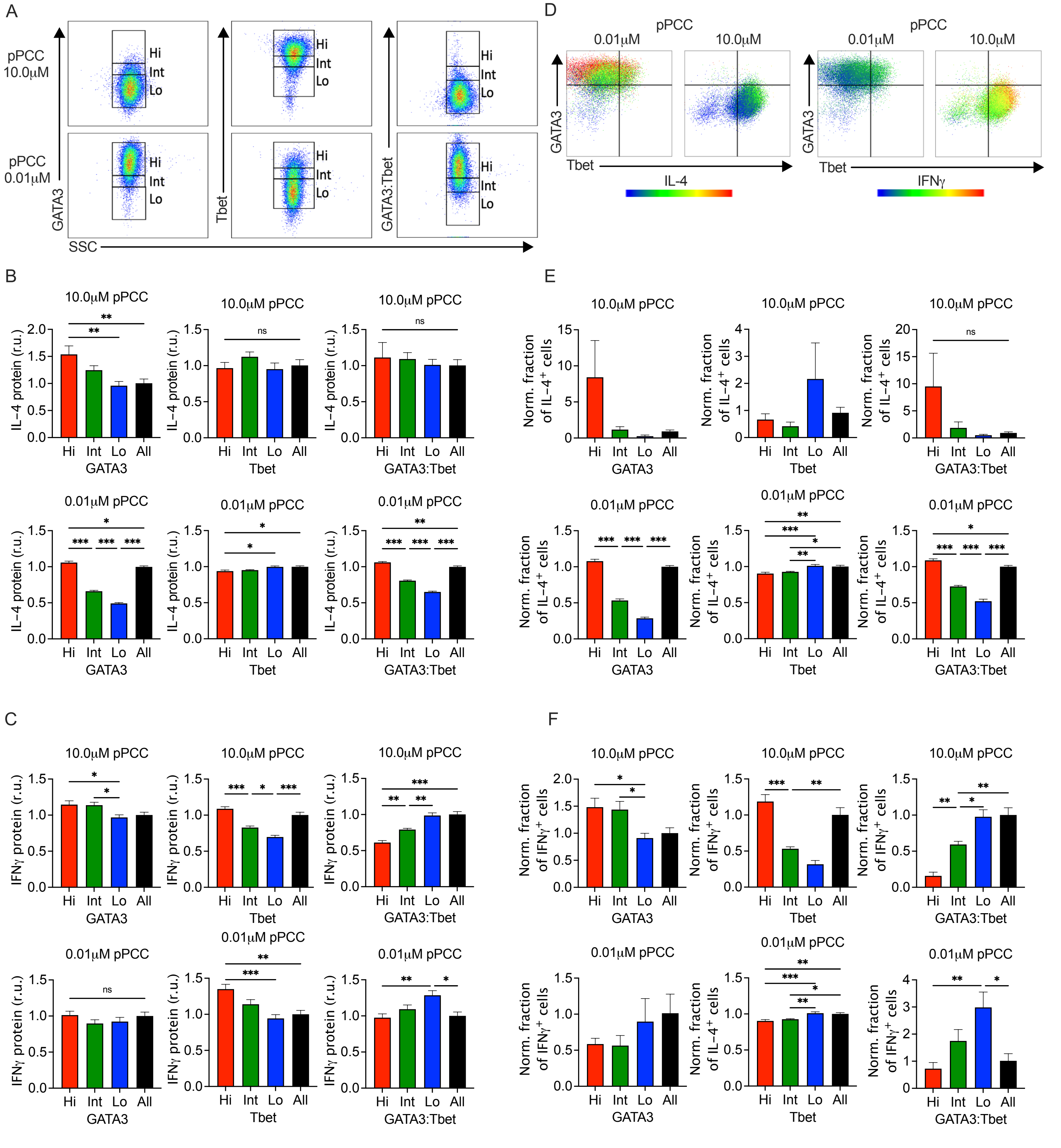

**Figure S7: Cytokine production and transcription factor expression are quantitatively correlated**

Naïve 5CC7 CD4+ T cells were stimulated with P13.9 antigen presenting cells in the presence of 0.01 µM or 10.0 µM pPCC for 4 days under in vitro conditions. (A) Following restimulation, the levels of GATA3, Tbet, and GATA3:Tbet expression were used to assign cells to high (Hi), intermediate (Int), or low (Lo) groups. Comparative analysis of IL-4 expression (B&E) and IFNγ expression (C&F) in cells grouped by relative level of transcription factor expression was conducted (All = relative units of MFI or % of total ungated CD4+ population). (D) Levels of IL-4 or IFNγ expression were heat-mapped to representative plots of GATA3/Tbet expression values for restimulated cells. (B,C,E, and F) Means are plotted ±SEM, n = 6. * = $p < 0.05$, ** = $p < 0.01$, *** = $p < 0.001$, ns = non-significant.

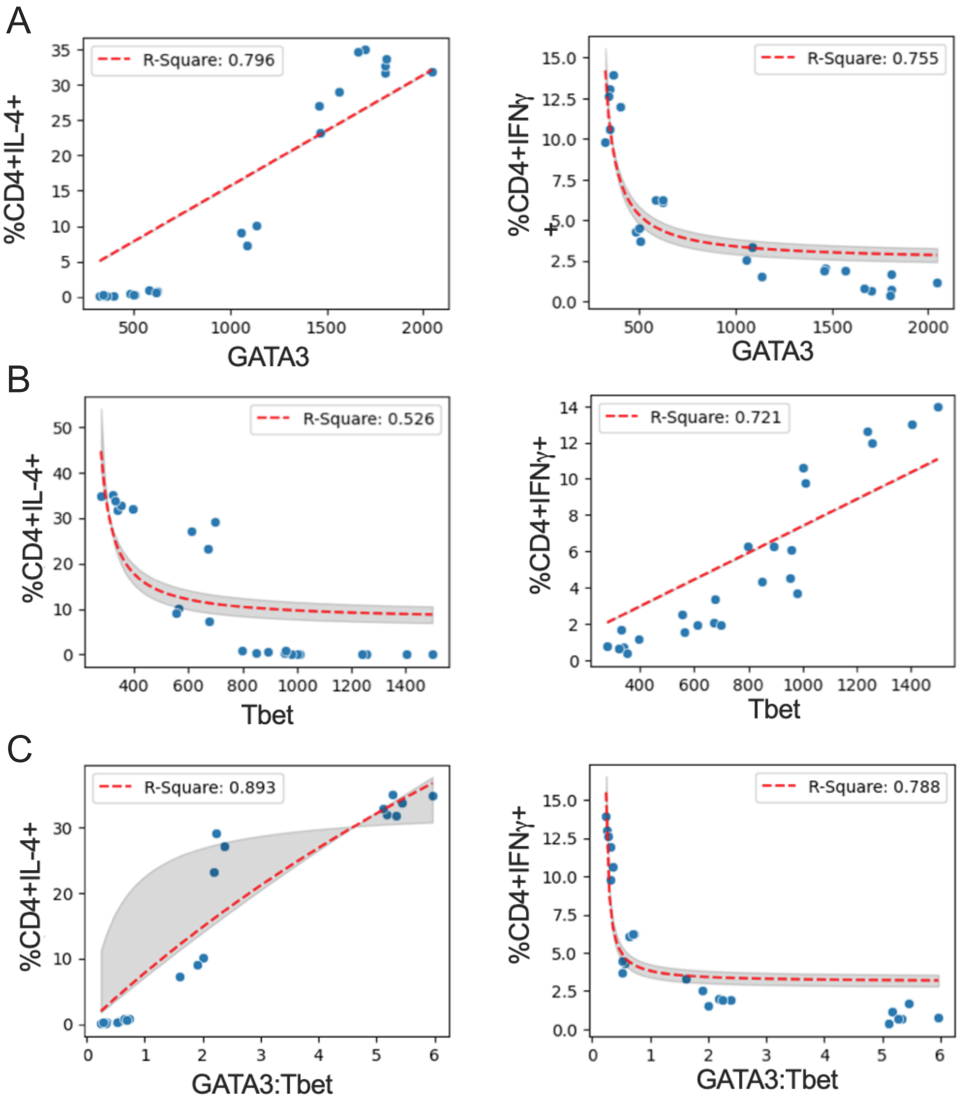


**Fig S8 Two-factor empirical dose response curve modelling indicates closest association between GATA3:Tbet and cytokine expression** Correlation between IFNγ+ or IL-4+ frequency and (A) GATA3, (B) Tbet, or (C) GATA3:Tbet expression from 5CC7 cells stimulated with 0.01 µM, 0.1 µM, 1.0 µM, or 10.0 µM pPCC. Means are plotted ±SEM, n = 6 per condition. The red line shows the best fit to the data obtained in model fitting conducted using a two-factor empirical dose response curve (Y = Ymax × (TF/K + TF), as in Helmstetter et al. [30]. The shaded region indicates the 95% confidence interval.

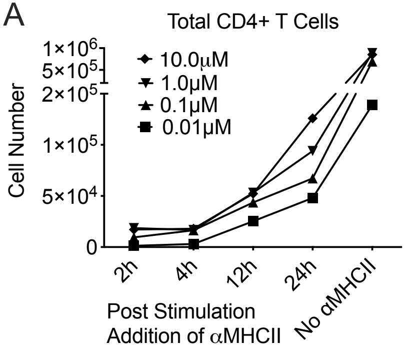


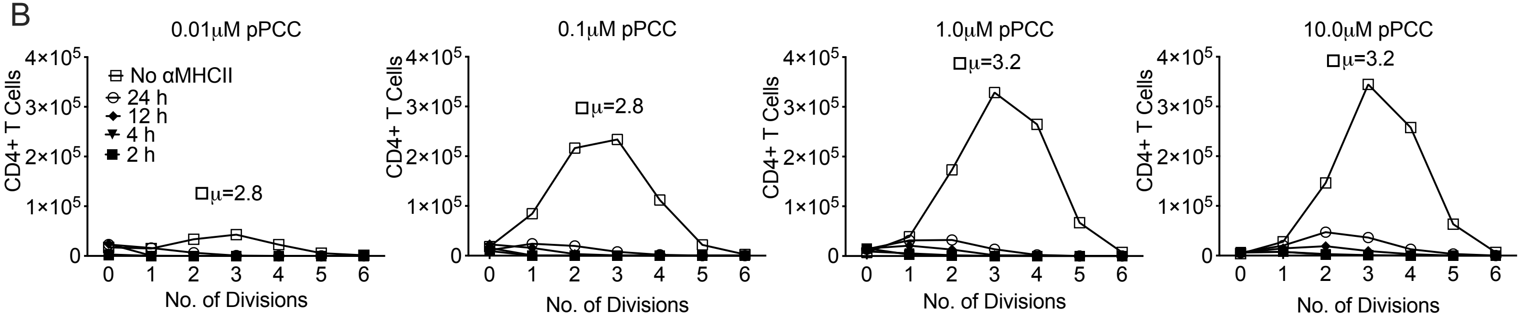


**Figure S9: CD4+ T cell expansion and survival are modulated by TCR signal strength and signal duration**

Naïve 5CC7 $CD4^+$ T cells were stimulated with P13.9 antigen-presenting cells in the presence of pPCC. CD4+ T cells were pre-labelled with CFSE to track division. Duration of TCR stimulation was controlled by the addition of anti-MHCII antibodies at the time points indicated to halt interactions. Induction of cellular division was assessed by CFSE dilution. (A) Total CD4+ cell counts were determined after stimulation. (B) Numbers of cells which had been through a specific number of cell divisions were calculated following analysis of CFSE dilution. All experiments were performed at least twice with consistent results. $\mu$ = mean number of divisions entered into by CD4+ T cells in the absence of anti-MHCII antibody.
